# Temporal phase-resolved transcriptomics reveals host determinants of Marek’s disease virus reactivation in transformed chicken T cells

**DOI:** 10.64898/2026.08.10.743901

**Authors:** Haji Akbar, Yung-Tien Tien, Kathrine Van Etten, Keith W. Jarosinski

## Abstract

Marek’s disease virus (MDV) is an oncogenic herpesvirus that establishes latency in CD4⁺ T cells, from which it reactivates to initiate productive replication and dissemination. To define host mechanisms governing the transition from latency to early and late lytic replication, we developed recombinant MDV expressing early RLORF4mRFP and late UL47eGFP, which was used to isolate latent (Lo; mRFP low), early lytic (Hi; mRFP high), and late lytic (DP; mRFP⁺eGFP⁺) populations from MDV-induced lymphoblastoid cell lines (LCLs). Two independent LCLs (Lines 62 and 82) were subjected to Illumina RNA sequencing after sorting-purified populations. Viral transcription increased progressively from Lo to Hi to DP populations; however, initiation of viral gene expression differed markedly between cell lines. During the early transition (Lo to Hi), Line 82 exhibited robust induction of 17 viral genes, whereas Line 62 showed only two differentially expressed viral genes. However, both lines converged on a highly conserved transcriptional program during the transition from latency (Lo) to late-lytic replication (DP), sharing 118 viral transcripts, most of which were structural and assembly-associated genes. Among the host transcriptional responses, Line 82 exhibited activation of TP53-associated stress signaling, chromatin remodeling factors, and RNA biogenesis pathways, consistent with a permissive cellular state. In contrast, Line 62 displayed enhanced inflammatory, metabolic, and proteostasis-associated signatures, suggesting a restrictive environment that limits early viral induction. Collectively, these findings demonstrate that MDV reactivation is a host-gated process in which the cellular state governs the initiation of the lytic switch, whereas downstream replication proceeds through a conserved viral program.

**IMPORTANCE:** This study provides critical insights into the host-virus dynamics governing herpesvirus reactivation from latency. By developing a dual-reporter system using recombinant Marek’s disease virus (MDV) in MDV-transformed lymphoblastoid cell lines (LCLs), an oncogenic herpesvirus model, reactivating cells were sorted for latent, early-lytic, and late-lytic populations from two independent LCLs. The initial transition from latency to early-lytic replication was highly variable and strongly influenced by cellular state: one line showed rapid viral gene induction linked to stress, chromatin remodeling, and permissive conditions, while the other exhibited a more restrictive, inflammatory/metabolic response. However, progression to late-lytic replication converged on a highly conserved viral transcriptional program dominated by structural genes. These findings establish that host cellular context gates the lytic switch in herpesvirus reactivation, offering a powerful framework for understanding latency control across oncogenic herpesviruses and potential therapeutic targets to prevent reactivation and tumor formation.

## INTRODUCTION

Marek’s disease is a lymphoproliferative disease of chickens caused by gallid alphaherpesvirus 2 (GaAHV2; species *Mardivirus gallidalpha2*), better known as Marek’s disease virus (MDV). MDV transforms chicken T lymphocytes and produces gross lymphomas in susceptible chickens, costing the poultry industry worldwide an estimated 1 billion US dollars annually (1). Although the International Committee on Taxonomy of Viruses (ICTV) classifies MDV in the *Alphaherpesvirinae* subfamily of the *Orthoherpesviridae*, the virus shares key biological attributes with members of the *Gammaherpesvirinae* subfamily. Specifically, MDV establishes latency in lymphocytes and transforms them into cancer. Tumor-derived cells can be cultured *ex vivo* to generate MDV-induced lymphoblastoid cell lines (LCLs). These LCLs provide a powerful experimental system to study herpesvirus-induced oncogenesis, latency, and reactivation (2). Thus, MDV infection and lymphocyte transformation in chickens offer a natural animal model for dissecting herpesvirus-induced lymphoid malignancy, herpesvirus reactivation, and host–virus interactions.

Latency constitutes a critical feature of herpesviruses. During latency, MDV integrates into host telomeres through MDV-encoded viral telomeric repeats (TMRs) (3). This integration enables stable maintenance of the viral genome during host cell division while simultaneously positioning the virus for efficient reactivation. During latency, herpesviruses evade immune recognition by limiting viral protein expression. They also suppress lytic viral gene expression and cellular translation by producing noncoding viral RNAs (4). Epigenetic modulation of the viral genome further regulates viral gene expression (5)—for example, by producing latency-associated transcripts (LATs) that negatively regulate MDV ICP4 and ICP27 (6). Reactivation initiates a lytic infection, which provides the virus with a survival strategy: it escapes from the latently infected cell before cell death, produces infectious progeny, and spreads to new cells or hosts.

The viral gene expression program during primary lytic replication for alphaherpesviruses such as herpes simplex virus (HSV) has been well established. The temporal cascade begins with immediate early (IE) genes, such as ICP4, ICP22, and ICP27, following viral entry into the cell. IE genes then induce the expression of early (E) genes, such as the thymidine kinase (TK) and conserved herpesvirus protein kinase (CHPK). After the initiation of viral DNA replication, late (L) genes encoding structural and envelope proteins are expressed. For MDV, unique genes such as pp38, pp14, and Meq also play crucial roles in regulating gene expression (7). However, it remains unclear whether lytic replication initiated during reactivation follows the same transcriptional hierarchy observed during primary infection. For gammaherpesviruses, the replication and transcription activator (Rta), encoded by ORF50, functions as the “switch” that initially activates expression of a multitude of cellular and viral genes (8). In contrast, investigators have not yet identified an equivalent viral “lytic switch” gene for MDV. We hypothesize that MDV reactivation may depend more heavily on host-cell–defined regulatory states, including chromatin accessibility, transcriptional competence, stress signaling, metabolic reprogramming, and immune modulation.

Several techniques to have been used to reactivate MDV in LCLs *ex vivo*, including hypoxia (9), serum starvation (10), chemical treatment (11), and temperature treatment (12). Temperature treatment induces reactivation of MDV in LCLs more effectively than sodium butyrate does, without causing significant increases in apoptosis or cell death. Hicks *et al.* (13) used microarray analysis to identify MDV-encoded microRNAs (miRNAs) during reactivation induced by sodium butyrate treatment. Similarly, Mwangi *et al.* (14) performed RNA-seq analyses of LCLs expressing fluorescently tagged late protein UL47 during spontaneous *ex vivo* reactivation. However, these studies largely focused on late lytic infection and did not fully resolve transcriptional changes associated with the transition from latency to early lytic activation. To address this critical knowledge gap, we employed two independent MDV-induced LCLs that express fluorescent reporter tags distinguishing the early and late phases of viral gene expression and replication (15–17). Using graded reactivation treatments, we captured transcriptional states spanning latency, early reactivation, and late lytic replication (Fig. 1). RNA sequencing coupled with systems-level pathway and upstream-regulator analyses was used to define host transcriptional programs governing MDV reactivation. This approach enabled us to identify shared and cell-line–specific host pathways that regulate permissiveness to lytic replication and to nominate candidate host regulators that control the MDV lytic switch.

**Fig. 1.**
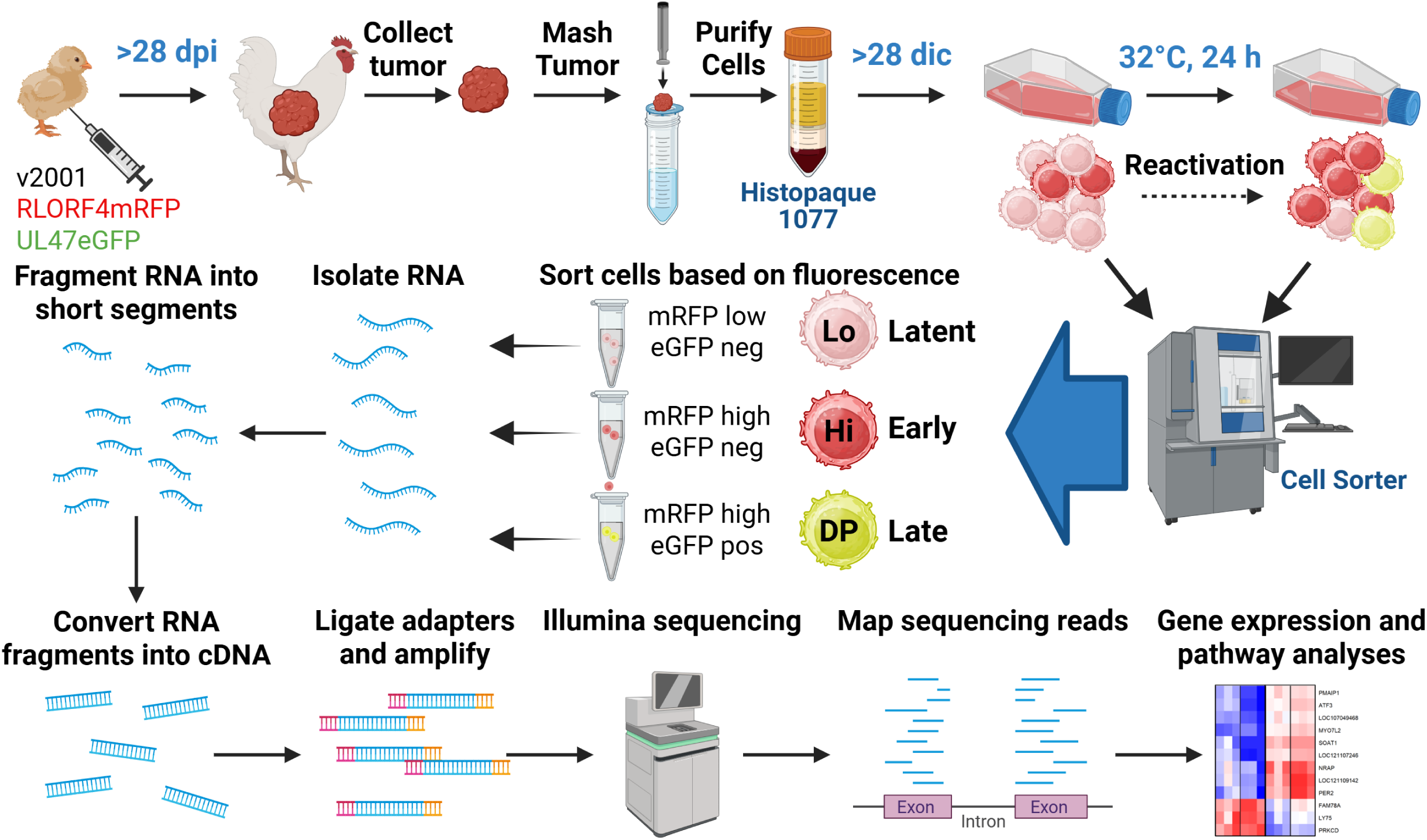
Experimental design for generating, characterizing, and sorting LCLs at different phases of replication following reactivation. Chickens were experimentally infected with a dual fluorescent virus (v2001) expressing RLORF4mRFP and UL47eGFP. Tumors were collected more than 28 days post-infection (dpi) and expanded for more than 28 days in culture (dic). Reactivation assays were used to reactivate MDV by treating cells at 32°C for 24 h. Cells with latent MDV express low levels of mRFP (Lo), while cells reactivating the virus express high levels of mRFP (Hi) during the early phase of replication but lack eGFP expression. Cells expressing mRFP and eGFP (double positive; DP) indicate that the virus has reached the late phase of replication. RNA-seq was performed on cell populations, and gene expression and pathway analyses were used to examine host cell responses to reactivation.

## RESULTS

### Generation and characterization of LCLs

Four independent MDV-transformed LCLs were established from v2001-infected chickens (17) and maintained *ex vivo* for >20 weeks. This virus expresses fluorescent reporters fused to the early protein RLORF4 (monomeric red fluorescent protein; mRFP) and the late protein UL47 (enhanced green fluorescent protein; eGFP), enabling the distinction of three populations: latent (mRFP low, Lo), early lytic (mRFP high, Hi), and late lytic (mRFP⁺eGFP⁺, double positive; DP). All LCLs were >99% CD4⁺ (Table S1), consistent with canonical MDV-induced T-cell phenotypes (18).

### Reactivation competency of LCLs

Low-temperature treatment (32°C for 24 h) induced reproducible reactivation in all lines, accompanied by increased expression of mRFP and eGFP (Fig. 2A & B). However, the magnitude of reactivation differed substantially between lines. Lines 62 and 82 exhibited the highest proportions of Hi and DP populations while maintaining high viability (Fig. 2B). In parallel, flow cytometric analysis revealed a decrease in CD4 expression in Hi and DP populations following reactivation (Fig. 2C), recapitulating known markers of MDV reactivation (14). Line 59 showed reduced viability following reactivation and was excluded from downstream analyses.

**Fig 2.**
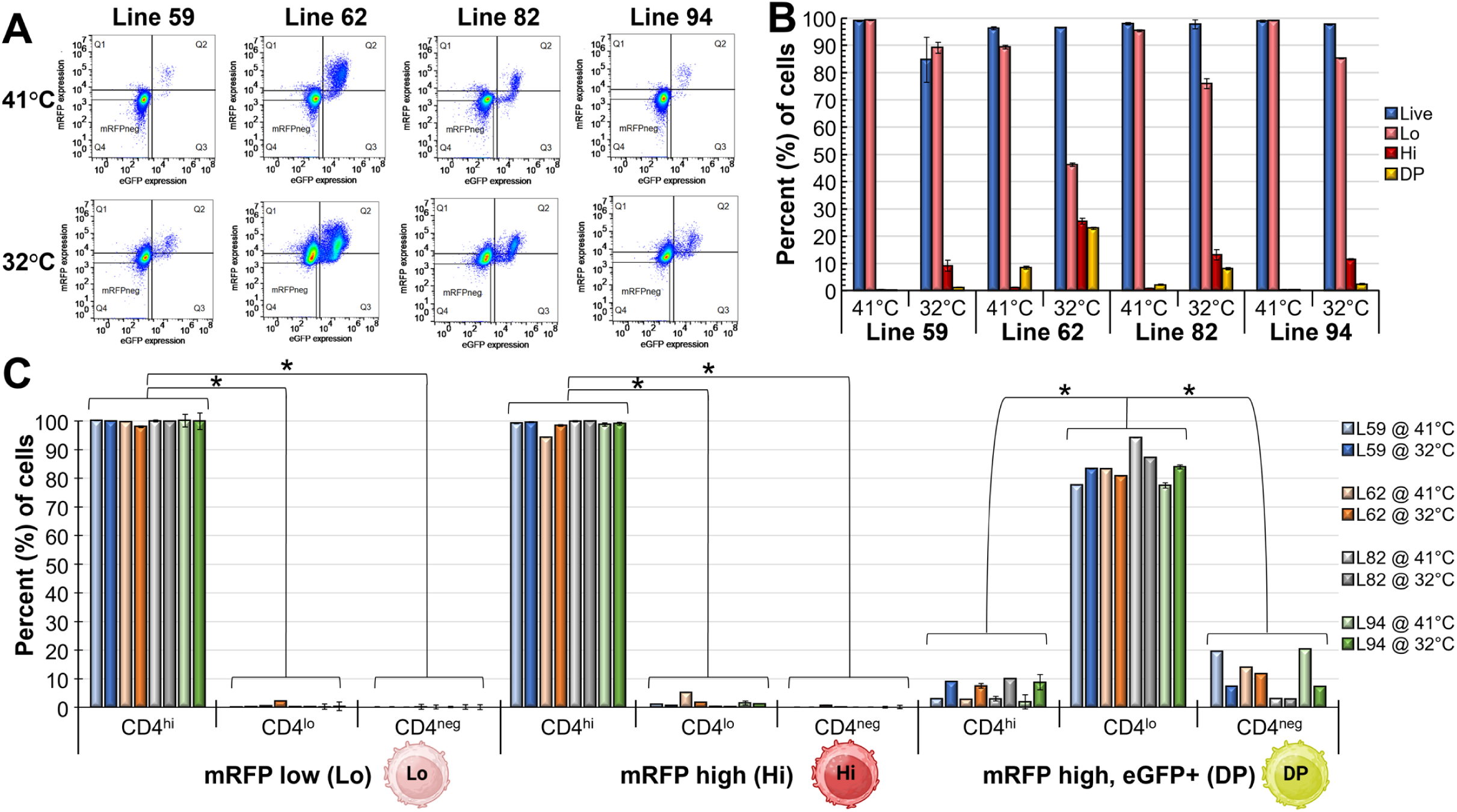
Characterization of MDCC lines. (A) Flow cytometry dot plots of mRFP and eGFP expression in LCLs. (B) Percentage of cells expressing mRFP low (Lo), mRFP high (Hi), and mRFP + eGFP (DP) with cell viability (Live). (C) CD4 surface expression on LCLs after low-temperature reactivation. Data in (B) and (C) are presented as mean ± standard deviation (n = 3). Statistical significance was determined using the Kruskal–Wallis test followed by Dunn’s multiple-comparison test with Bonferroni adjustment, as indicated by (*; P < 0.05).

### Cell-free virus production following reactivation

Next, we examined the ability of each cell line to produce cell-free virus following reactivation through temperature treatment (32°C for 24 h) and collection of cell-free virus extracts. Plaque-formation assays demonstrated significant increases in cell-free virion production following temperature-induced reactivation, with Line 82 producing the highest titers (Fig. 3), indicating greater permissiveness for productive replication.

**Fig 3.**
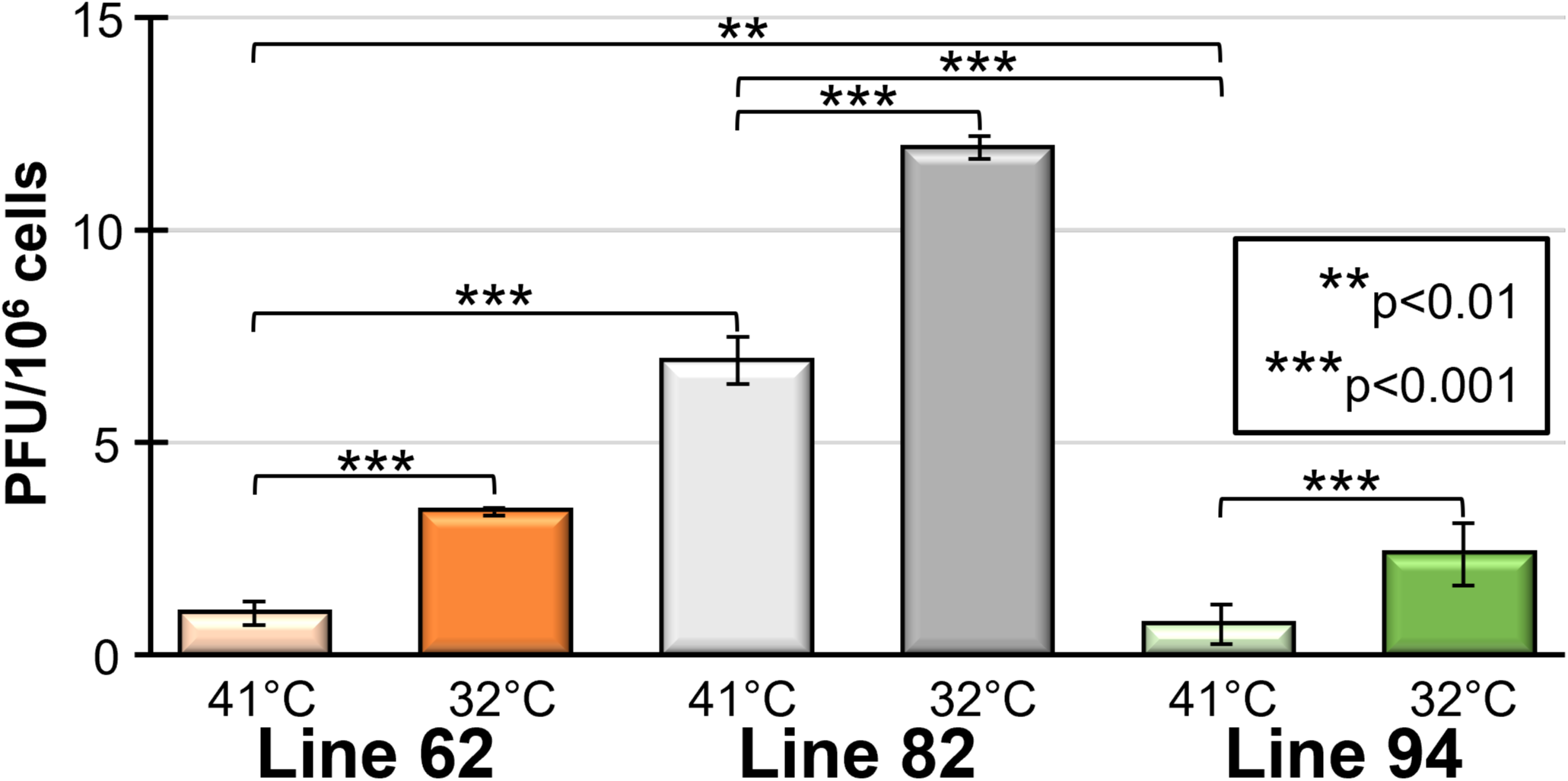
Reactivation potential of LCLs. Plaque-forming units for Lines 62, 82, and 94 following temperature treatment (32°C) to induce reactivation, followed by cell-free virus extraction and plaque-formation assays. Statistical significance was determined using the Kruskal–Wallis test followed by Dunn’s multiple-comparison test with Bonferroni adjustment (*; P < 0.05).

Overall, these data show that independent LCLs vary significantly in their reactivation potential, and fluorescent protein expression in LCLs transformed by v2001 is a robust indicator of MDV reactivation and progression through the early and late replication stages. Lines 62 and 82 showed the highest proportions of Hi and DP populations and cell-free viruses, while maintaining high viability; therefore, they were selected for transcriptomics analysis.

### Viral gene expression following reactivation

Lo, Hi, and DP populations from Lines 62 and 82 (n =3 per group) were subjected to RNA-seq analysis (Fig. 1). Approximately 1 billion clean reads were generated, averaging 55.8 million reads per sample. Viral transcript abundance increased progressively from Lo to Hi to DP populations in both lines (Fig. 4, Fig. S1A & B), validating stage-specific capture of reactivation progression. Low levels of viral transcripts were detected in the Lo populations of both LCLs, confirming basal viral transcription during latency. Viral reads increased stepwise from Lo to Hi to DP in both LCLs, validating stage-resolved capture of the lytic transition. Overall viral transcript abundance in the Lo population was greater in Line 62 than in Line 82 (Fig. 4, Fig. S1A & B).

**Fig 4.**
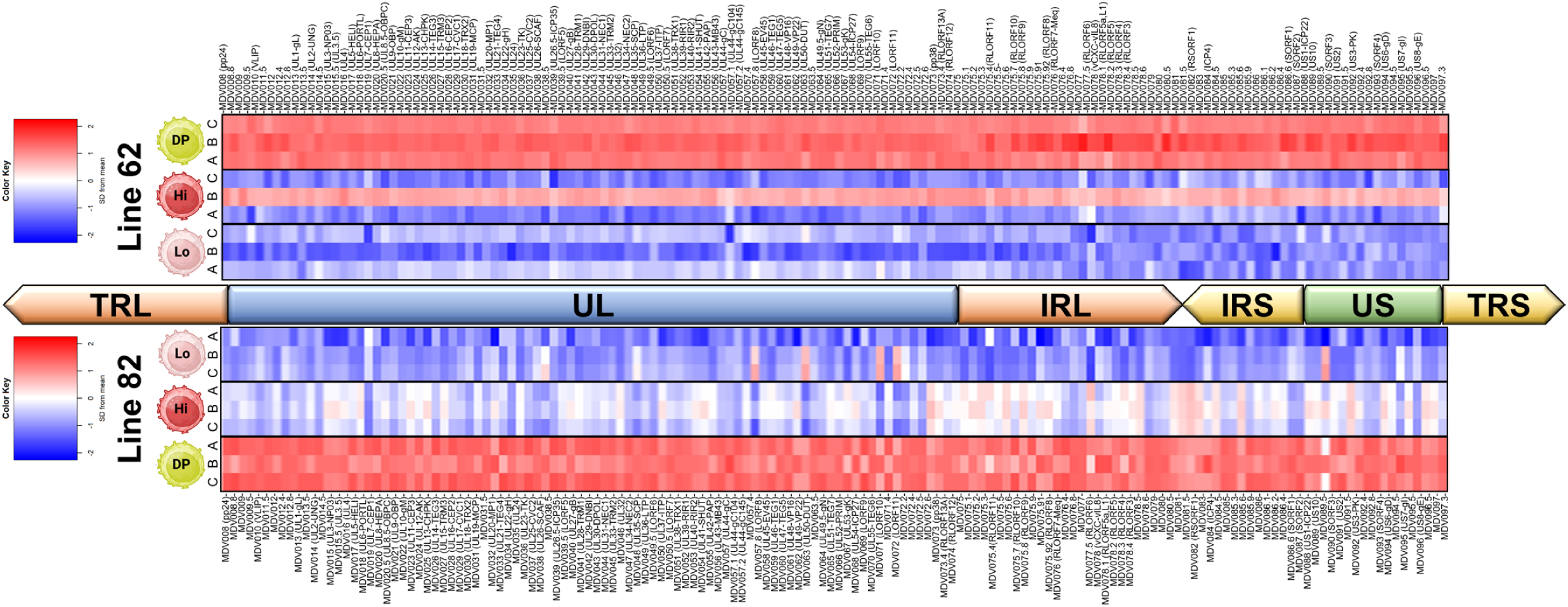
Heatmap of MDV gene expression following reactivation. A heatmap of gene expression for all MDV genes from latent (Lo), early lytic replication (Hi), and late lytic replication (DP) groups is shown. Shown are the unique long (UL) and short (US) and internal repeat long (IRL) and short (IRS) used in the alignment. The terminal repeat long (TRL) and short (TRS) regions were trimmed from the alignment.

### Viral gene expression during latency to early lytic replication

During the transition from latency (Lo) to early lytic (Hi) replication, differential expression was highly cell-line-dependent: Line 82 exhibited 17 viral differentially expressed genes (DEGs), whereas Line 62 showed only 2, with no genes shared between the LCLs at FDR < 0.05 (Data S1). Early-induced viral genes in Line 82 included *MDV019* (*UL7*), *MDV028* (*UL16*), *MDV047* (*UL34*), *MDV065* (*UL51*), and *MDV092* (*US3*), together with several RLORF-associated transcripts.

### Viral gene expression from latency to late lytic replication

In contrast, transitions involving the DP population revealed highly conserved viral transcriptional programs between the two cell lines (Fig. 4, Data S1). From latency (Lo) to late lytic replication(DP), 140 and 120 DEGs were identified in Lines 62 and 82, respectively, with 118 shared transcripts. Similarly, the early (Hi) to late (DP) lytic comparison revealed 133 viral DEGs in Line 62 and 89 in Line 82, with 88 genes shared. These common transcripts were dominated by structural and assembly genes characteristic of productive herpesvirus replication, including *MDV023* (*UL11*), *MDV040* (*UL27*), *MDV048* (*UL35*), *MDV050* (*UL37*), *MDV051* (*UL38*), *MDV060* (*UL47*), and *MDV064* (*UL49.5*), defining a conserved late lytic cascade independent of LCL.

These results show an expected transition in viral gene expression from latency (Lo) to early lytic replication (Hi), followed by late lytic replication (DP). However, our results also show that viral gene expression and reactivation potential differ considerably between Lines 62 and 82. Together, these findings indicate that initiation of viral transcription is likely cell-line dependent, whereas the downstream late lytic program is highly conserved once productive replication is established.

### Transcriptomic profiling reveals cell line- and replication stage-specific gene expression fates

Multidimensional scaling (19) of the 5,000 most variable genes showed separation by both cell line and replication stage (Fig. S1C), indicating that host transcriptional trajectories differ between LCLs despite identical reactivation conditions. DEGs were analyzed using the limma-voom method (19) with a model of Group + 5 RUV factors for both cell lines. Due to differences between the two LCLs, three pairwise comparisons were performed within each cell line: Hi vs. Lo, DP vs. Lo, and DP vs. Hi (Fig. 5, Table 1, Data S1). When examined separately, both cell lines showed distinct gene expression patterns (Fig. 5). For example, 3 and 505 DEGs were identified in Lines 62 and 82, respectively, when comparing the Hi and Lo groups (Fig. 5A, Table 1, Data S1). The DP vs Hi comparison revealed 20 and 780 DEGs (FDR < 0.05, FC >2) in Lines 62 and 82, respectively, with seven genes shared between both LCLs (Fig. 5B, Table 1, Data S1). In the DP vs Lo comparison (Fig. 5C, Table 1, Data S1), 85 and 1,220 DEGs (FDR < 0.05, FC > 2) were identified in Lines 62 and 82, respectively, with 37 genes shared between the two LCLs (Table 1, Data S1). Shared DEGs included genes associated with proteostasis (HSPA2, DNAJB1, HSPB9), immune signaling modulation (NFKBIE, SOCS3, DUSP8), transcriptional regulation (TP53, ASH1L), and cytoskeletal/adhesion processes (MARCKSL1, EPHA2, ITGA4). These 37 common host genes likely constitute a conserved host program required for productive MDV replication.

**Fig 5.**
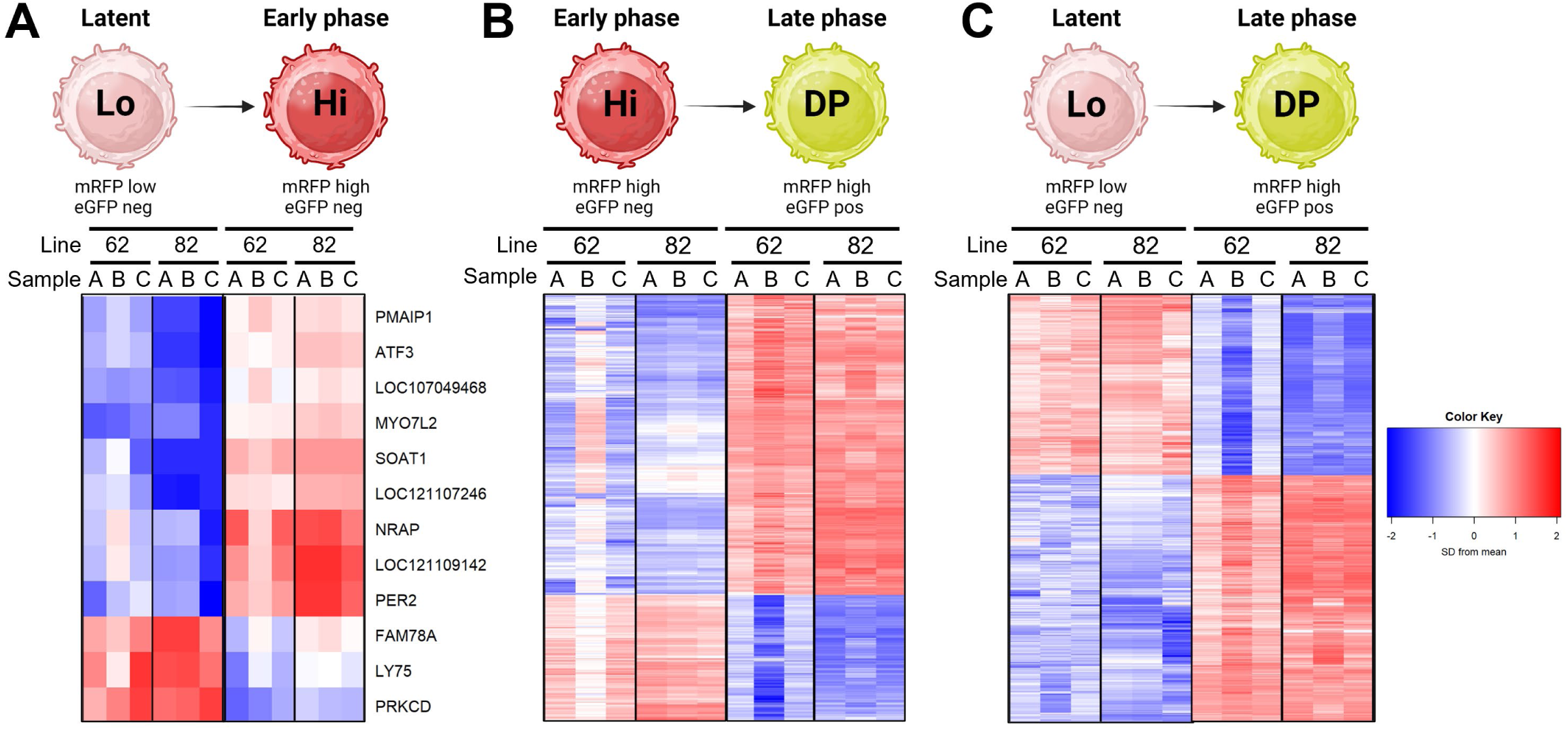
Heatmaps comparing Line 62 and Line 62 during the transition from latent to early and late lytic replication. (A) Heatmap of gene expression for 12 genes showed a significant, consistent direction of change when comparing early replication (Hi) to late (Lo) cells in both cell lines. (B) Heatmap of 853 genes with the same direction of change in late (DP) compared to early (Hi) phase replication in both lines. (C) Heatmap of 437 genes showing changed expression in the same direction as the change in late phase (DP) compared to latent (Lo) cells in both lines.

**Table 1.** The number of differentially expressed genes (global FDR p-value < 0.05) and the top 10 up- and down-regulated genes within each of the 6 pairwise comparisons.

| Comparison | Line | DEGs<br>(Down/Up) | Shared<br>(Down/Up) | Top Ten (Down/Up) DEGs |
| --- | --- | --- | --- | --- |
| Hi vs Lo | 62 | 3<br>(1/2) | None | AACS<br>LOC121107013, LOC771494 |
|  | 82 | 505<br>(119/386) | None | NPTN, LOC769729, DNAJC12,<br>LOC101747645, BLB2, PCASP2,<br>HOPX, HEXDC, LOC112532284,<br>LOC107051359<br>LOC121107204, LOC121108097,<br>LOC107054459, ACTG1, C2H5ORF22,<br>SPNS2, C30H19orf53, KAT2A, ISCA1,<br>SLC25A25 |
| DP vs Hi | 62 | 20<br>(1/19) | None | HP1BP3<br>LOC107054772, LOC121107472,<br>LOC112530097, LOC121107331,<br>LOC121106809, LOC121113400,<br>C19orf25, LOC121107343, KLF4,<br>LOC121106791 |
|  | 82 | 780<br>(389/391) | 7<br>(0/7) | LOC112531610, P2RY8,<br>LOC121106450, CDH11, PLEK,<br>PM20D1, ZMYND8, TFB1M,<br>PRTFDC1, TTC7B<br>LOC395991, HSPB9, RORBL,<br>LOC112532562, LOC124417094,<br>LOC121108930, LOC121111700,<br>LOC121107276, MARCKSL1, GLRA4 |
| DP vs Lo | 62 | 85<br>(9/76) | None | LOC100859645, ITGA4, SMARCA2,<br>ARFGAP3, IDE, IQGAP3, RAP1A,<br>PLAGL1, ARHGDIA,<br>LOC112530097, LOC121106560,<br>LOC112532707, LOC107054772,<br>LOC101750589, LOC121108930,<br>LOC121107472, NFKBIE,<br>LOC121108910, C19orf25 |
|  | 82 | 1220<br>(470/750) | 37<br>(2/35) | CDH11, PROKR1, LOC420160, SLA,<br>CD38, LOC769729, LOC121113176,<br>UNC13D, GPR1, MCTP1<br>HSPB9, LOC121108930,<br>LOC112530392, LOC121107243,<br>LOC121107383, LOC107049468,<br>EPHA2, LOC112530219,<br>LOC112532220, LOC121111251 |

### Enrichment analysis overview

The striking divergence in reactivation between Lines 62 and 82 suggested that initiation of the MDV transcriptional cascade may be governed by host-cell permissiveness, consistent with the model that Line 62 maintains a more restrictive state during the earliest step of reactivation, whereas Line 82 is less restrictive. To identify potential cellular responses to these differences, differential gene expression results were interrogated using Ingenuity Pathway Analysis (IPA, http://www.ingenuity.com/; Ingenuity Systems Inc., Redwood City, CA) and SRplot (https://www.bioinformatics.com.cn), an online platform for gene ontology (GO) and pathway enrichment visualization (20). At a stringent differential expression threshold (FDR < 0.05), IPA revealed a clear divergence in pathway-level inference between the two LCLs, mirroring the magnitude of differences observed at the gene level (Fig. 6A). Line 82 generated robust enrichment of transcription, antiviral responses, antigen presentation, protein homeostasis, “state-switching”, and classic herpesvirus late-phase replication control (cell cycle and DNA replication). Both cell lines had interactions among core viral, cell fate, trafficking, disease, and biofunctions (Fig. 6B); however, Lines 62 and 82 responded differently during the viral replication phase transitions. For example, Line 62 generally exhibited inhibitory responses from early during reactivation (Hi v Lo), followed by activation during transition from early to late (DP v Hi) lytic replication, while Line 82 had activation of pathways during early reactivation (Hi v Lo) that were inhibited during transition from early to late (DP v Hi) lytic replication. However, both cell lines primarily exhibited inhibitory responses, comparing latent to late (DP v Lo). This pattern supports a model of cell-line–dependent transcriptional permissiveness rather than experimental inconsistency, since both cell lines ultimately responded similarly, despite differences in the transition to late-lytic replication.

**Fig 6.**
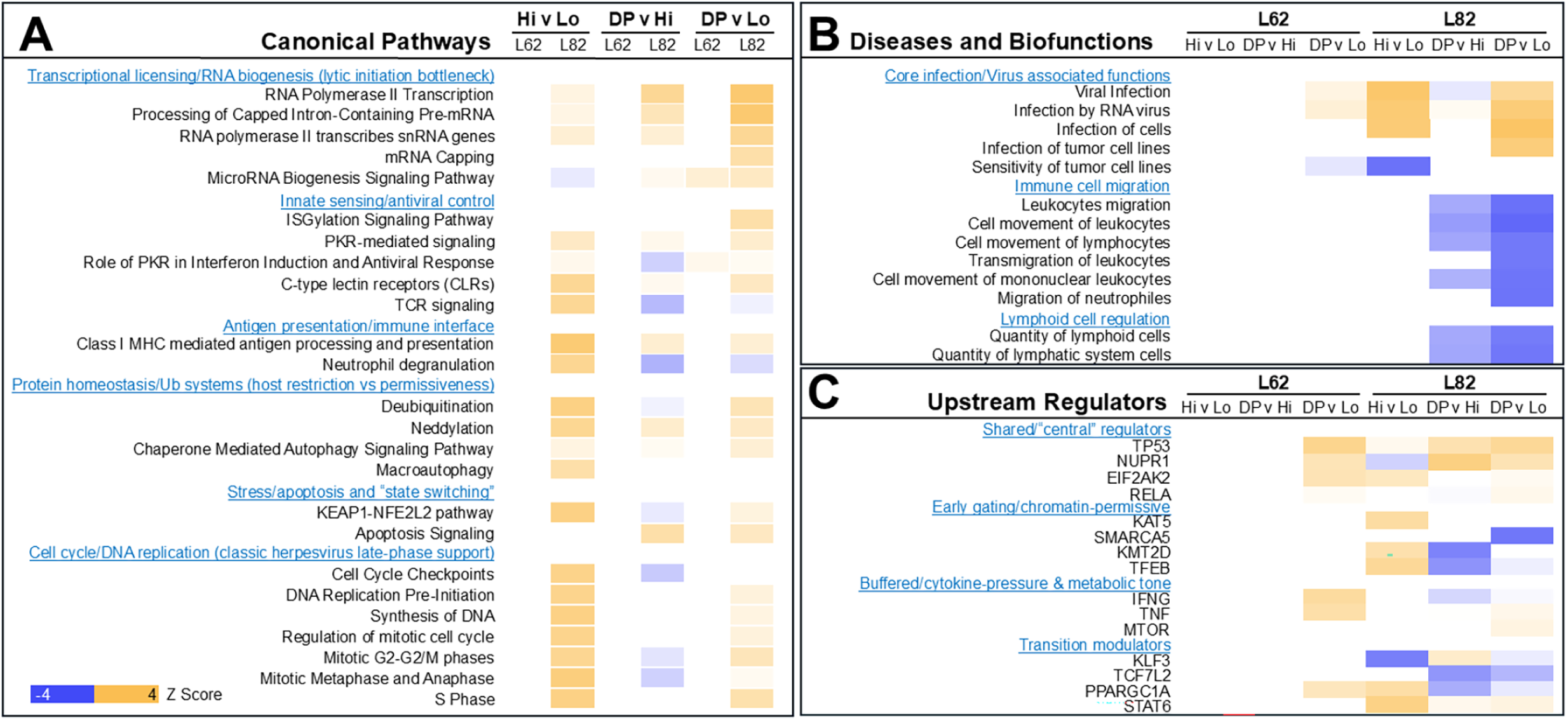
Integrated pathway, regulator, and functional enrichment analysis of host responses during MDV reactivation. IPA of RNA-seq data identified the top significant Canonical Pathways (A), Diseases and Bio Functions (B), and Upstream Regulators (C) associated with transitions from latency (Lo) to early (Hi) and late (DP) lytic replication in MDV-infected lymphoblastoid cell lines (Line 62; L62 and Line 82; L82). Results represent a combined analysis across both cell lines and all stage comparisons (Hi vs. Lo, DP vs. Hi, and DP vs. Lo), with full cell line–specific results provided in Data S2. The color reflects the predicted activation state based on the IPA z-score (blue: inhibition; orange: activation). Differential expression input thresholds were FDR < 0.05, and enrichment significance was defined by IPA at p-value < 0.05.

Despite the lower number of DEGs in Line 62, both LCLs showed concordant enrichment of core virus-response functions at FDR < 0.05. Disease and biofunction analysis identified shared activation of the categories “Viral Infection” and “Infection by RNA virus” in late (DP) vs latent (Lo) for both cell lines, with stronger activation scores in Line 82 (Fig. 6B). These common signatures demonstrate that both LCLs engage bona fide antiviral and infection-associated programs during reactivation, albeit with different amplitudes.

Because Line 62 showed fewer DEGs at an FDR < 0.05, a relaxed threshold (FDR < 0.25) was applied for pathway-level discovery, an accepted strategy to capture coordinated but lower-magnitude transcriptional programs (21). This approach enabled the recovery of coherent biological signals in Line 62 while retaining the high-confidence framework established for Line 82 (Fig. 7). Applying the FDR < 0.25 threshold substantially expanded the detectable transcriptional landscape in both LCLs, most notably in Line 62, where the number of DEGs and pathway coverage increased while retaining directional concordance with the stringent analysis. The late (DP) vs latent (Lo) comparison—representing the most complete transition from latency to productive replication—remained the dominant contrast in both lines, supporting its designation as the principal molecular representation of the MDV lytic switch.

**Fig 7.**
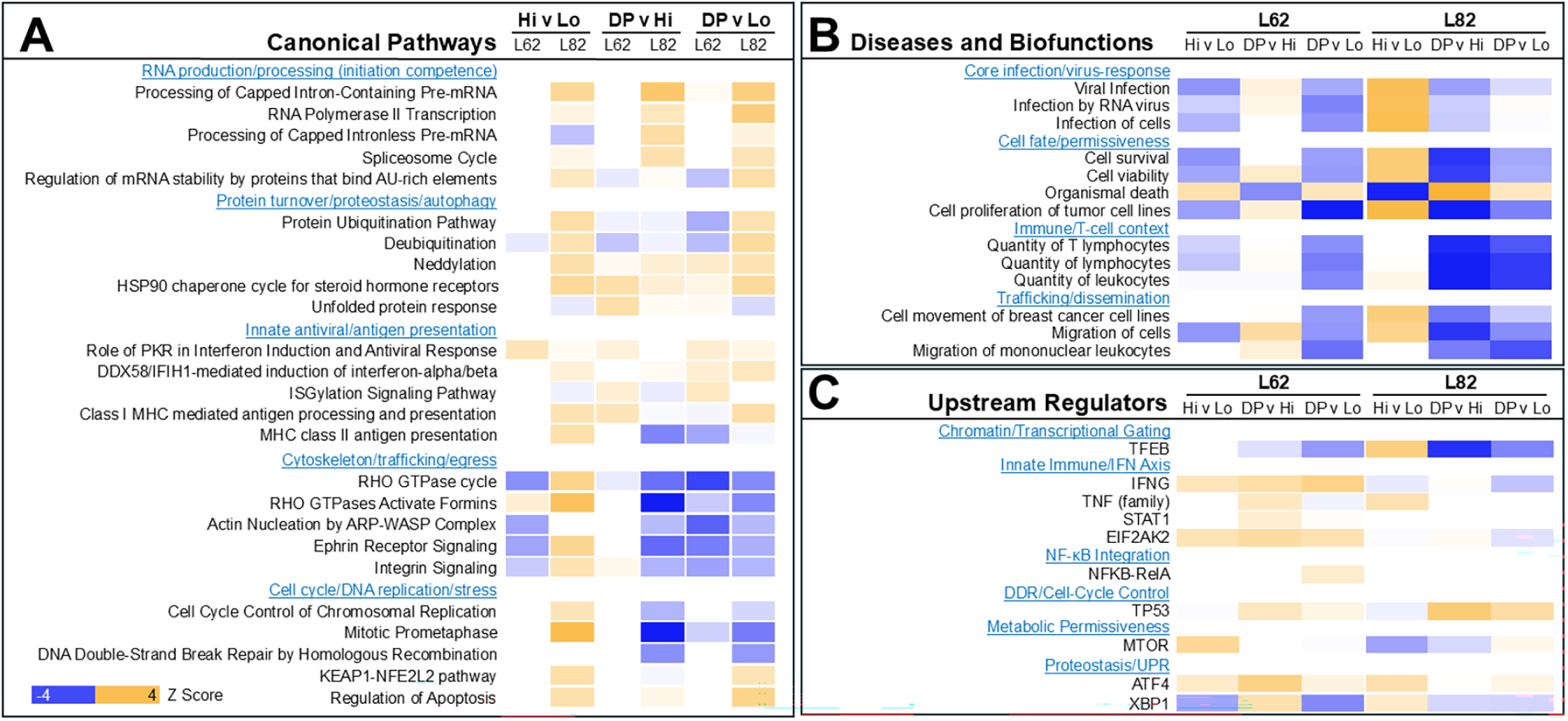
Integrated pathway, upstream regulator, and functional enrichment analysis of host responses during MDV lytic reactivation. IPA of RNA-seq data identified the top significant Canonical Pathways (A), Diseases and Bio Functions (B), and Upstream Regulators (C) associated with transitions from latency (Lo) to early (Hi) and late (DP) lytic replication in MDV-infected lymphoblastoid cell lines (Line 62; L62 and Line 82; L82). Results represent a combined analysis across both cell lines and all stage comparisons (Hi vs. Lo, DP vs. Hi, and DP vs. Lo), with full cell line–specific results provided in Data S2. The color reflects the predicted activation state based on the IPA z-score (blue: inhibition; orange: activation). Differential expression input thresholds were FDR <0.25, and enrichment significance was defined by IPA at p-value < 0.05.

### Transcriptional licensing associated with early reactivation in Line 82

During the transition from latency (Lo) to early (Hi) lytic replication (FDR < 0.05), pathway inference was largely restricted to Line 82, consistent with the minimal number of DEGs observed in Line 62 (Fig. 6). In Line 82, early reactivation was associated with activation of host transcriptional licensing and RNA biogenesis pathways, innate sensing, antigen presentation, protein homestasis, the stress response, and classical herpevirus late-phase replication support (Fig. 6A).

### Limited transcriptional licensing during early reactivation in Line 62

In contrast to Line 82, Line 62 exhibited limited pathway enrichment during transitions from latent (Lo) to early lytic replication (Hi) using the high stringency (FDR < 0.05); however, relaxing the FDR to 0.25 provided more information (Fig. 7A). Interestingly, Line 62 showed inhibition of cytoskeletal traffiking, while Line 82 showed activation of this pathway among others during early reactivation (Hi v Lo).

### GO and KEGG pathway enrichment analyses

To capture coordinated systems-level transcriptional programs that may not meet stringent statistical thresholds, GO (Fig. 8A & B) and KEGG (Fig. 9A & B) pathway enrichment analyses were performed using SRplot on DEGs identified with FDR < 0.25.

**Fig. 8.**
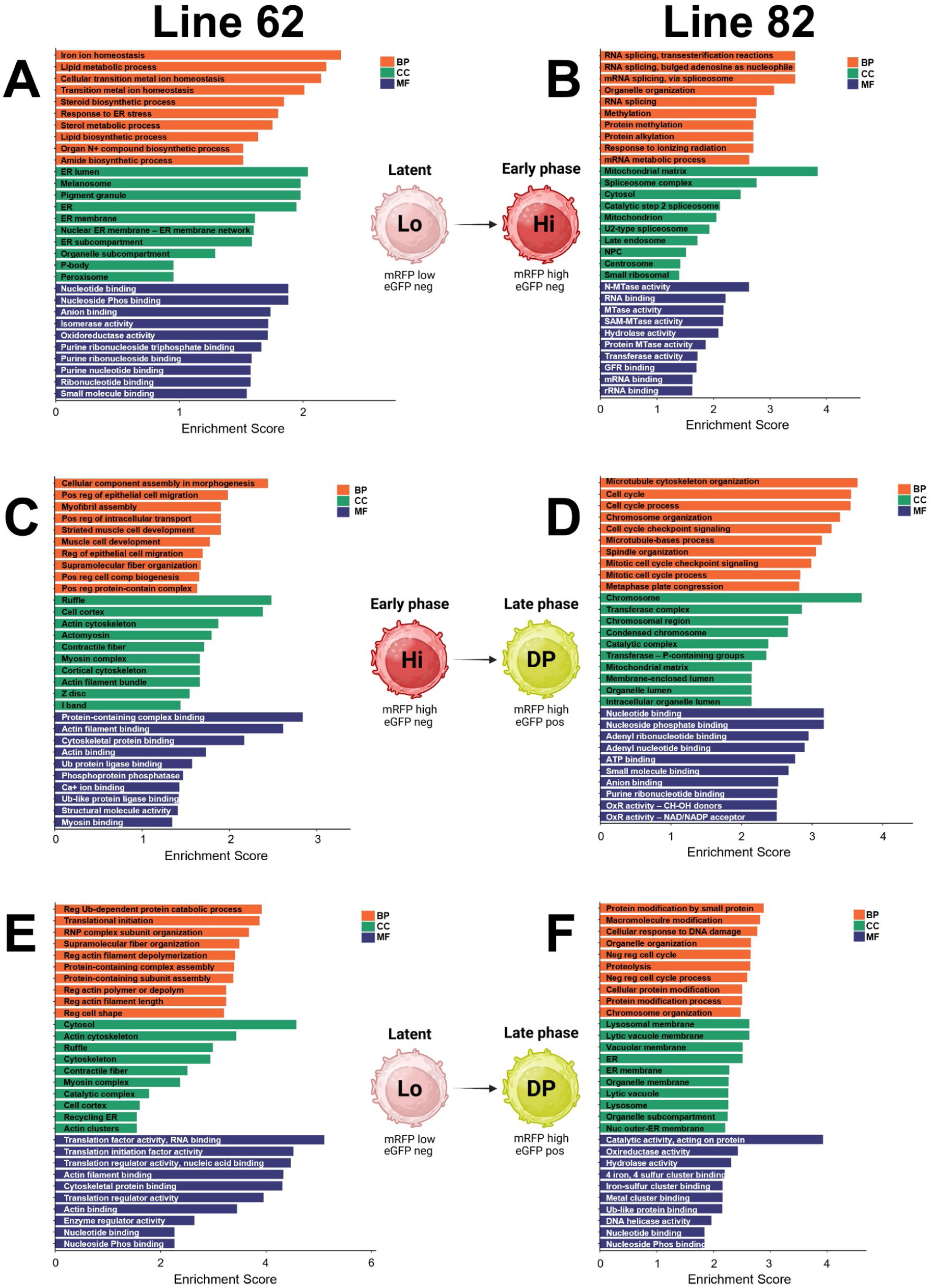
Gene Ontology (GO) enrichment analysis of host cellular responses during MDV reactivation in LCLS. GO enrichment analysis was performed using SRplot on differentially expressed genes identified with FDR < 0.25 for each stage transition and cell line. Panels A, C, and E represent Line 62, and panels B, D, and F represent Line 82. Cells are shown transitioning from latent (Lo) to early lytic (Hi) replication (A-B), from early (Hi) to late lytic (DP) replication (C-D), and from latent (Lo) to late lytic (DP) replication (E-F). Bars represent significantly enriched GO terms across Biological Process (BP), Cellular Component (CC), and Molecular Function (MF) categories. The length of each bar reflects the enrichment score (–log10 adjusted p-value), indicating the relative statistical significance of each term. These results demonstrate stage-dependent and cell-line–specific activation of host processes involved in transcriptional regulation, protein synthesis, intracellular transport, metabolic remodeling, and stress and immune responses during MDV reactivation. Complete cell–line–specific results are provided in Data S2.

**Fig. 9.**
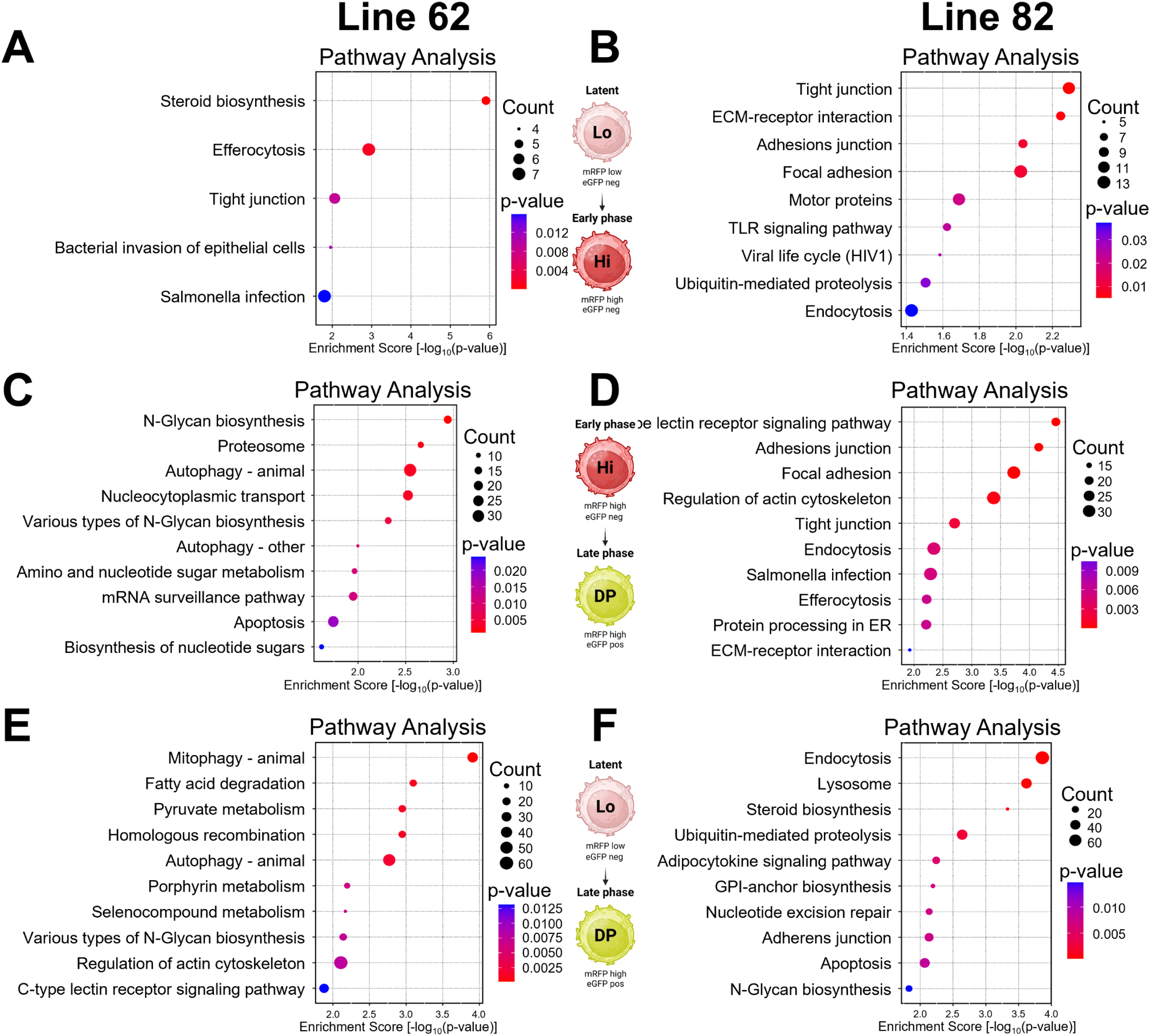
KEGG pathway enrichment analysis of host cellular programs during MDV reactivation in LCLs. Pathway enrichment analysis was performed using SRplot on differentially expressed genes identified with FDR < 0.25 for each stage transition and cell line. Panels A, C, and E represent Line 62, and panels B, D, and F represent Line 82. Representation of cells transitioning from latent (Lo) to early lytic (Hi) replication (A-B), from early (Hi) to late lytic (DP) replication (C-D), and from latent (Lo) to late lytic (DP) replication (E-F). Each dot represents a significantly enriched pathway, with dot size indicating the number of genes mapped to the pathway and color representing enrichment significance (adjusted p-value). These results reveal progressive engagement of host transcriptional, metabolic, and stress-response pathways during MDV reactivation, with stronger pathway activation observed during the late lytic phase and in the more permissive Line 82. Complete cell line–specific results are provided in Data S2.

### Transition from latency to early replication is host-gated

In Line 62, relaxed-threshold analyses primarily identified pathways associated with lipid metabolism, steroid biosynthesis, ER membrane organization, and translational regulation, with comparatively weak enrichment of RNA-processing pathways (Figs. 8A & 9A). In Line 82, GO enrichment analysis of DEGs (FDR < 0.25) revealed a strong representation of RNA processing and metabolic processes, including RNA splicing, mRNA metabolic process, spliceosomal complex assembly, and nucleocytoplasmic transport (Fig. 8B). KEGG pathway analysis showed strong representation of cell interaction and signaling pathways (Fig. 9B). Together, these data indicate that early MDV reactivation is associated with robust transcriptional and RNA-processing activation in the more permissive Line 82, whereas Line 62 shows comparatively limited host remodeling during the initiation of viral transcription.

### Early to late transition is associated with structural, biosynthetic, and trafficking

The early (Hi) to late (DP) lytic replication transition highlighted further activation of transcriptional processes, while other pathways activated early during reactivation (Hi v Lo) were inhibited in transitioning from early to late viral replication (DP v Lo) in Line 82 (Fig. 6A). For example, innate sensing/viral control, antigen presentation, herpesvirus late-phase support were activated from latency to early lytic replication (Hi v Lo), but inhibited from early to late lytic (DP v Lo). Comparisons between early and late lytic replication (DP v Hi) at relaxed-threshold analysis (FDR < 0.25) identified host remodeling associated with progression toward productive late-stage replication. For Line 62, a similar but less robust response was evident, in which pathways activated during reactivation (Hi v Lo) were inhibited when transitioning from early to late-phase replication (DP v Hi).

GO pathway analyses identified enrichment in cellular morphogenesis in Line 62 (Fig. 8C), while Line 82 showed enrichment for mRNA cell-cycle processing and surveillance, nucleic acid binding, and cytoskeleton rearrangements for Line 82 (Fig. 8D). However, both lines displayed overlapping enrichment of pathways involved in intracellular trafficking, protein processing, and structural remodeling. KEGG pathway analysis showed enrichment for autophagy and biosynthesis of sugar in Line 62 (Fig. 9C) and cell adhesion and tight junctions in Line 82 (Fig. 9D). In all, these changes coincided with cellular changes promoting viral gene transcription, translation, and cytoskeletal rearrangements, supporting coordinated host remodeling during productive MDV replication.

### Coordinated host remodeling is associated with fully productive lytic replication

The overall transition from latency to fully productive replication (DP v Lo) comparison captured the most robust and biologically coherent signatures in both LCLs, especially Line 82 (Figs. 6A & 7A). IPA identified coordinated enrichment of pathways associated with RNA production and processing, protein turnover, cell-cycle progression, stress responses, and innate antiviral signaling during latency (Lo) to late lytic (DP) replication (Fig. 7A). In Line 82, highly enriched activated canonical pathways included RNA production and processing, protein turnover and proteostatsis, and innate antiviral responses, while cytoskeletal signaling and cell trafficking and cell cycle and DNA replication were inhibited (Fig. S2). Line 62 also showed enrichment for inhibition of cytoskeletal signaling and cell trafficking, suggesting this is a consistent response during fully productive reactivation.

Using the relaxed-threshold analyses (FDR < 0.25) further expanded pathway coverage, particularly in Line 62, while retaining directional concordance with the stringent analysis. In Line 62, GO enrichment emphasized cytoskeletal organization and regulation of actin polymerization (Fig. 8E). In Line 82, enriched KEGG pathways included ubiquitin-mediated proteolysis, lysosome, endocytosis, apoptosis, nucleotide excision repair, and glycan biosynthesis (Fig. 9F). Although enrichment magnitude differed between lines, both exhibited coordinated remodeling of cytoskeletal and proteostasis pathways during late replication.

### Candidate host determinants associated with MDV permissiveness

Integration of canonical pathways and network analyses identified several host upstream regulatory nodes consistently associated with stage-specific MDV reactivation. Across analyses, TP53, MTOR, XBP1, NFKB1/RELA, IFNG/TNF-associated signaling, EIF2AK2, DDX58, and STUB1/BAG1-centered proteostasis modules emerged as prominent regulators linked to differences between the two LCLs (Figs. 6C & 7C). Line 82 consistently exhibited stronger activation of transcriptional, stress-response, and chromatin-associated pathways, whereas Line 62 displayed comparatively greater enrichment of inflammatory, proteostasis, and ER stress–associated signaling. Collectively, these analyses identify distinct host transcriptional states associated with permissive versus restricted initiation of MDV reactivation.

### Coordinated activation of pathways associated with MDV reactivation

Disease and biofunction analyses identified strong enrichment for viral infection in both cell lines at FDR < 0.05, although the magnitude of enrichment was consistently greater in Line 82 (Fig. 6B). At an FDR <0.25, both cell lines had consistent enrichment for overall activation of core responses for viral infection, cell fate, T cell immunity, and trafficking. Although similar functional categories were detected in both LCLs, Line 82 exhibited stronger enrichment of transcriptional and replication-associated functions, whereas Line 62 showed relatively greater enrichment of stress- and immune-related processes.

## DISCUSSION

Herpesviruses maintain infection throughout the life of their host. Reactivation from latency is an important part of the herpesvirus life cycle, providing an opportunity to disseminate widely throughout the population, and it remains one of the least understood stages. Previous MDV transcriptomic studies have compared latent and late cytolytic replication phases (13, 14); however, herpesvirus gene expression occurs in a temporal cascade, progressing from immediate-early to early to late phases of infection. In this study, we examined the transformed T cells’ response to herpesvirus replication across three phases: latency, early lytic replication after reactivation, and fully productive replication, using MDV-transformed LCLs expressing fluorescent reporters to separate latent (Lo: mRFP low), early lytic (Hi: mRFP high), and late lytic (DP: mRFP high plus eGFP-positive or double-positive: DP) populations (Fig. 1). Four independent LCLs were established, each derived from a different chicken. Following further characterization for reporter expression, CD4 down-modulation, and cell-free virion production, it was clear that each LCL differed in its ability to reactivate MDV, consistent with the concept of tumor heterogeneity (22). To further address viral gene expression and host cell response to reactivation, we selected Lines 62 and 82 for transcriptomic analysis (Fig. 2). Line 82 exhibited substantially greater viral gene expression and broader host transcriptional remodeling, whereas Line 62 displayed a buffered response, indicating intrinsic differences in permissiveness rather than technical variation.

Our data confirm the uniqueness of individual cancers and, despite their induction of transformation by similar factors (MDV infection). Using this system, clear conserved mechanisms or pathways are involved, whereas each LCL’s response during reactivation differs.

### Host-dependent initiation of MDV reactivation

The earliest stage of MDV reactivation (Lo to Hi) showed the greatest divergence between the two LCLs. Following temperature-induced reactivation, Line 82 showed induction of 17 viral transcripts with FDR < 0.05, whereas Line 62 showed only 2, with no shared viral DEGs between the lines comparing early lytic replication (Hi) from latency (Lo) (Fig. 4, Data S1). Low levels of viral transcripts were detected in the latent (Lo) populations of both LCLs, with overall viral transcript abundance being greater in Line 62 than in Line 82. This observation suggests that the two LCLs may differ in the transcriptional state associated with latency or in the mechanisms regulating entry into productive reactivation, although this hypothesis requires further experimental validation. Line 62 showed only minimal induction of viral genes during the Lo-to-Hi transition, possibly due to the higher basal abundance of viral transcripts in the latent (Lo) population. This may indicate that successful initiation of productive reactivation depends on host cellular permissiveness rather than on basal viral transcript abundance alone. However, both cell lines expressed virtually all viral transcripts when transitioned to late-gene expression, as indicated by the expression of mRFP and eGFP (DP). Because both lines were exposed to identical induction conditions and were sorted using the same reporter logic (RLORF4mRFP for early and UL47eGFP for late), these results argue that the critical barrier lies in the host state that permits early viral transcription rather than in differences in the downstream viral program. In other herpesviruses, lytic initiation is known to be limited by chromatin structure and transcriptional licensing at viral promoters, and modest differences in host permissiveness can translate into large differences in immediate-early/early viral transcript output (23–26).

### Conserved and divergent host responses during MDV reactivation

Our use of two enrichment thresholds served complementary purposes. The stringent (FDR < 0.05) IPA analyses identify high-confidence pathway and regulator signals tightly linked to robust gene-level changes, whereas the relaxed (FDR < 0.25) analyses—used for systems-level discovery in both IPA and SRplot—recover coordinated modules that may reflect subtle but biologically meaningful shifts, especially in less responsive contexts such as Line 62. This two-tier strategy is increasingly used in transcriptomic and reactivation studies, where initiating changes may be modest, distributed across regulatory networks, yet decisive for phenotypic outcomes (21). Importantly, conclusions drawn from FDR < 0.25 are framed here as hypothesis-generating and are supported by concordant signals across multiple enrichment tools (IPA and SRplot) and by parallel viral transcriptional progression.

Across both cell lines, canonical pathway analysis revealed coherent modules consistently modulated relative to latently infected cells (Lo) and recently reactivated cells (Hi), compared with late viral gene expression or fully productive replication (DP) (Figs. 7 & S2). Prominent among these were: (i) cytoskeletal and trafficking remodeling, including inhibition of Rho-GTPase–related pathways, consistent with reorganization required for virion assembly and egress; (ii) host translation and RNA production machinery, with suppression of eukaryotic translation initiation signaling and enrichment of RNA processing terms; and (iii) lipid and cholesterol metabolic regulation, reflected by coordinated inhibition of multiple cholesterol biosynthesis pathways. These convergent signatures indicate that both LCLs engage in fundamental cellular processes that are exploited during herpesvirus lytic replication, despite differences in transcriptional amplitude. By combining stage-resolved sorting (Lo, Hi, DP) with parallel viral and host transcriptomics in two independent MDV-transformed LCLs (Lines 62 and 82), our data support a biphasic model of the lytic switch: (i) an initiation step that is strongly host-gated and cell-line dependent, followed by (ii) execution of a largely conserved late viral transcriptional cascade once the DP state is reached. This framework aligns with general herpesvirus biology in which chromatin accessibility, transcription/RNA-processing capacity, stress responses, and innate immune tone collectively shape lytic competence (23, 24, 27–29). Thus, MDV latency and reactivation are tightly regulated processes in which a small number of host “gatekeeper” programs determine whether cells remain in a latent/transformed state or transition into productive (lytic) replication.

### Host transcriptional permissiveness determines early lytic activation

At the host gene transcriptional level, Line 82 exhibited substantially greater host and viral transcriptional responses than Line 62 at stringent thresholds (FDR < 0.05), particularly during the transition from latency (Lo) to early replication (Hi). This suggests that reactivation is not merely a passive consequence of the temperature shift but reflects cell-line–specific permissiveness. Importantly, this “amplitude difference” was echoed at the pathway level. For example, IPA identified enrichment of robust canonical pathways, upstream regulator predictions, and disease and biofunction networks in Line 82, whereas Line 62 yielded fewer—but mechanistically coherent—signals, most evident compare latent (Lo) to late replication (DP) phase cells (Fig. 6). These observations are consistent with prior reports that clonally derived MDV LCLs display divergent reactivation phenotypes and reinforce the value of analyzing multiple independent lines when defining host determinants of reactivation (12–14, 30).

Consistent with this interpretation, early lytic replication-stage host pathway signals in Line 82 were dominated by RNA biogenesis and RNA fate, processes essential for productive herpesvirus gene expression (Fig. 6). Canonical pathway enrichment at FDR < 0.05 highlighted RNA Polymerase II transcription, capped intron pre-mRNA processing/splicing, mRNA capping, and regulation of mRNA stability, together indicating a transcriptionally competent and post-transcriptionally permissive environment. Independent GO and KEGG enrichment using SRplot on relaxed-threshold DEGs (FDR < 0.25) further reinforced these themes in Line 82 (e.g., RNA processing, spliceosome-related terms, nucleocytoplasmic transport, and mRNA surveillance), suggesting that reactivation is coupled to broad remodeling of host RNA processing capacity rather than to a single regulatory node (Fig. 7B). Herpesviruses are highly dependent on host RNA polymerase II, splicing/processing machinery, and mRNA export, and several viral proteins directly manipulate these steps to favor viral transcript maturation and translation (23, 31–33).

In contrast, Line 62 showed a comparatively muted host response transitioning from latency to early lytic replication (Lo-to-Hi), with limited pathway detection at FDR < 0.05, but a distinct enrichment profile at FDR < 0.25 that was skewed toward lipid/steroid metabolic processes and ER-associated functions (Fig. 9A). This divergence suggests that Line 62 may require additional host “licensing” events to enter a fully transcriptionally permissive state, consistent with the established role of host chromatin remodeling, transcriptional activation, and RNA-processing machinery in enabling herpesvirus reactivation (24, 26, 34). Interestingly, overall viral transcription early in lytic replication (Lo) was greater in Line 62 than in Line 82, suggesting that the two LCLs differ in the transcriptional state associated with latency or in the mechanisms regulating entry into productive reactivation. This observation is consistent with the idea that herpesvirus latency is a dynamic continuum rather than a strictly transcriptionally silent state (14, 35–37). However, despite the higher basal viral transcript abundance observed in Line 62, this cell line exhibited only minimal induction of viral genes during the Lo to Hi transition, indicating that basal viral transcript abundance alone is insufficient to drive productive reactivation and further supporting the conclusion that host cellular permissiveness governs entry into the lytic program. Alternatively, the muted transcriptional shift from latency (Lo) to early lytic replication (Hi) in Line 62 suggests that temperature induction elicits only partial stress and metabolic responses, without fully engaging the nuclear transcriptional expansion necessary for efficient viral gene expression — consistent with stress signals being insufficient to trigger full herpesvirus reactivation in the absence of coordinated transcriptional licensing pathways (37–39). Together, these findings suggest that Line 62 exists in a transcriptionally buffered host state that restricts early viral gene induction following temperature treatment, whereas Line 82 more readily transitions into a transcriptionally permissive configuration that enables efficient initiation of the lytic cascade. This buffered phenotype is further supported by reduced pathway recovery at stringent thresholds and predominance of stress, lipid, and ER-associated programs at relaxed thresholds.

### Host remodeling supports productive late-lytic replication

During progression to late replication (Lo vs DP and Hi vs DP), host remodeling converged on biosynthetic, proteostatic, and structural programs that matched the requirements of virion production. In Line 82, IPA at FDR < 0.05 revealed coordinated enrichment of protein turnover pathways (deubiquitination and neddylation), cell-cycle and DNA replication modules (S-phase/checkpoint signaling), oxidative stress and apoptosis-related pathways (KEAP1–NFE2L2/Nrf2), and innate antiviral modification (ISGylation) (Fig. 6). These findings are consistent with the known dependence of herpesviruses on host ubiquitin–proteasome and proteostasis systems for viral protein maturation, capsid assembly, and immune evasion (40, 41), as well as recruitment of host DNA replication and repair machinery to facilitate viral genome amplification (42, 43). Activation of oxidative stress and Nrf2 pathways further reflects host metabolic and stress adaptation during productive infection (44–46), while ISGylation represents a key antiviral modification system engaged during herpesvirus replication (47–49). Using SRplot at FDR < 0.25 similarly highlighted ubiquitin-mediated proteolysis, lysosome/endocytosis, apoptosis, and DNA repair, comparing latent (Lo) to late-phase lytic (DP) replication (Fig. 8), consistent with increased viral protein synthesis, intracellular trafficking, capsid assembly, and nuclear egress that accompany productive alphaherpesvirus replication (50–55).

### Disease and biofunction enrichment analyses support the transition toward productive MDV replication

IPA revealed strong enrichment in categories associated with core virus infection and responses, trafficking/dissemination, and immune function in the context of T cells (Fig. 7). These signatures likely reflect the increasing biosynthetic demand required for large-scale viral transcript production, tegument protein synthesis, capsid assembly, and virion maturation during progression into late lytic replication. Similar shifts toward enhanced transcriptional output and protein-processing capacity have been reported during productive replication of other herpesviruses, in which host biosynthetic machinery is extensively repurposed to support viral amplification (56–58). In parallel, enrichment of cellular stress response, inflammatory response, and immune signaling functions corresponded with upstream regulator predictions involving TP53, IFNG, TNF, and NF-κB pathways (Figs. 6-9), suggesting simultaneous activation of host defense mechanisms and stress-adaptation programs during reactivation. Although these functional signatures were detected in both LCLs, Line 82 consistently exhibited stronger enrichment of transcriptional and replication-associated functions, whereas Line 62 retained comparatively greater representation of immune and stress-related processes. This distinction further supports the concept that host cellular state influences permissiveness to MDV reactivation, while conserved biosynthetic and replication programs dominate once productive lytic replication is established. Consistent with this interpretation, the extensive overlap of late viral structural genes among LCLs suggests that host-dependent regulation primarily governs reactivation initiation, whereas downstream replication follows a comparatively conserved, virus-driven program.

The network structure provided additional mechanistic insight into why Line 62 appears “buffered.” Whereas Line 82 formed multiple high-scoring networks enriched for inflammatory/antimicrobial responses, cell-cycle control, and autophagy/interaction modules, Line 62 produced fewer networks dominated by chaperone–ubiquitin and proteostasis components, including BAG1, heat shock proteins, and the ubiquitin ligase STUB1/CHIP, as well as the innate sensor DDX58 (RIG-I) (Data S2). The CHIP/STUB1–HSP chaperone system is a central regulator of protein quality control and stress adaptation, balancing folding capacity and ubiquitin-mediated degradation of misfolded proteins (59–61). Viruses rely heavily on host proteostasis networks for proper folding of structural and tegument proteins, and chaperone availability can directly influence viral replication efficiency (62, 63). A proteostasis-centered architecture could therefore restrict early lytic progression by prioritizing protein quality control and stress buffering over large-scale transcriptional expansion. Consistent with this interpretation, the unfolded protein response (UPR) and ER stress pathways are known regulators of herpesvirus lytic permissiveness, in which activation of XBP1, ATF4, and related stress pathways can either promote or restrain lytic reactivation, depending on timing and magnitude (64–66). In parallel, DDX58 (RIG-I) functions as a key innate immune sensor that detects viral RNA and activates antiviral transcriptional programs that restrict herpesvirus replication (67, 68). Together, these findings suggest that Line 62 maintains a proteostasis- and stress-buffered cellular state that may limit transcriptional permissiveness and delay entry into the full lytic cascade.

### Host regulatory networks governing lytic permissiveness

At an FDR < 0.05, Line 82 prominently implicated chromatin and transcriptional regulators (e.g., KAT5/TIP60, SMARCA5, KMT2D) together with stress and autophagy regulators (e.g., ATF4/DDIT3 and TFEB) and the shared master regulator TP53 (Fig. 6C). These factors represent plausible host determinants of lytic competence because chromatin remodeling and histone modification directly regulate accessibility of viral promoters and enable the transcriptional burst required for lytic initiation (69–71). Chromatin remodelers and histone modifiers, including TIP60 and SWI/SNF-family complexes, regulate viral transcription by altering nucleosome positioning and epigenetic accessibility at viral genomes (72, 73). In parallel, TP53-linked stress and checkpoint programs, frequently activated during herpesvirus infection, coordinate DNA damage signaling, cell-cycle control, and metabolic remodeling that facilitate viral genome replication and late gene expression (74–76). Stress-responsive transcription factors such as ATF4 and DDIT3, along with the lysosomal regulator TFEB, further influence viral replication by regulating autophagy and proteostasis pathways required for efficient virion assembly and replication (77–81). Together, these upstream regulators form a mechanistic framework in which chromatin accessibility, stress signaling, and autophagy-associated remodeling serve as host-gating nodes that control entry into productive MDV replication.

At the relaxed threshold (FDR < 0.25), upstream regulator analysis expanded the regulatory landscape—particularly for Line 62—and revealed divergences that provide testable mechanistic hypotheses (Fig. 7C). Line 62 showed stronger predicted inflammatory and interferon-associated cytokine pressure (e.g., IFNG, TNF-family regulators, and STAT1/IRF-linked signaling), together with differences in MTOR and XBP1-driven unfolded protein response (UPR) balance relative to Line 82. These regulatory axes are well positioned to gate permissiveness, as interferon-STAT1 and NF-κB signaling pathways are known to suppress or reshape herpesvirus lytic transcriptional programs and maintain latency or restrict replication (82–84). MTOR signaling integrates metabolic and nutrient cues to regulate translation, protein synthesis, and biosynthetic capacity required for viral replication (85–87), while XBP1-dependent UPR activation directly influences herpesvirus lytic transcription and replication by modulating ER capacity and stress adaptation pathways (64, 88–90). Conversely, Line 82 exhibited stronger TP53-linked stress and cell-cycle remodeling together with broader engagement of RNA-processing and transcriptional machinery, processes that are critical for efficient viral gene expression and productive replication (39, 75, 76, 91). These findings support a model in which inflammatory and proteostatic signaling biases in Line 62 maintain a more restrictive cellular environment, whereas stress-coupled transcriptional licensing in Line 82 enables more efficient initiation of the MDV lytic cascade.

Host regulatory factors control initiation, whereas viral replication follows a conserved program Importantly, the viral transcriptome data mirror the host regulatory architecture discussed. Although early viral induction was strongly cell-line-dependent (17 vs 2 DE viral genes from Lo to Hi), the late viral transcriptional program was highly conserved, with the majority of DE viral genes shared between lines in Lo to DP (118 shared genes) and Hi to DP (88 shared genes). These shared genes were dominated by structural and assembly components, including UL27 (gB), UL35 (VP26), UL37 (ITP), UL47 (TEG5), and UL49 (VP22), consistent with the conserved late transcriptional cascade characteristic of herpesvirus replication (36, 92–94). This conservation strongly supports a model in which host determinants primarily regulate the initiation of reactivation, whereas once a critical permissiveness threshold is reached, the downstream replication program proceeds through a largely virus-encoded, stereotypic cascade that is relatively insensitive to host background (29, 95). Consequently, host regulatory factors are most likely to influence early initiation events (Lo-to-Hi transition) and cellular progression into productive replication states, rather than late structural gene expression itself.

Several early-induced viral genes in Line 82 further support the concept that reactivation requires rapid mobilization of virion assembly and nuclear egress machinery. These included UL7 (cytoplasmic envelope protein (CEP) 1, CEP1), UL16 (CEP2), UL34 (nuclear egress complex (NEC); NEC2), UL51 (TEG7), and US3 (protein kinase), which are known to regulate capsid maturation, nuclear egress, tegument assembly, and cytoskeletal remodeling during herpesvirus replication (96–103). The early expression of these genes during reactivation suggests that MDV may rapidly prepare cellular structural and trafficking systems to support virion assembly and transport. This differs from primary infection, where entry and genome delivery represent early bottlenecks, whereas during reactivation in already infected T cells, the primary requirement is re-establishing productive assembly and nuclear egress. Herpesviruses are known to extensively reorganize host cytoskeletal and nuclear transport systems to facilitate virion maturation and intracellular trafficking, supporting this interpretation (32, 104, 105)

### Candidate host determinants of MDV reactivation

Beyond pathway-level inference, the shared host DEG sets between lines during latency (Lo) and early virus replication (Hi) to productive replication (DP) provide concrete candidates for functional follow-up. Shared genes mapped to proteostasis (HSPA2, DNAJB1, HSPB9), immune signaling modulation (NFKBIE, SOCS3, DUSP8), transcriptional and epigenetic regulation (TP53, ASH1L, ZC3H11A-like), and cytoskeletal and adhesion trafficking (MARCKSL1, EPHA2, ITGA4) (Data S2). These functional categories align closely with established cellular requirements for productive herpesvirus replication. Herpesviruses impose a substantial burden on host protein-folding and quality-control machinery due to high levels of viral structural protein synthesis, thereby making heat shock proteins and co-chaperones essential for efficient virion assembly and stability (106–108). Concurrently, herpesviruses actively modulate host immune signaling pathways, including NF-κB and SOCS-regulated cytokine signaling, to balance the suppression of antiviral defenses with the maintenance of cell viability (109, 110). Transcriptional and chromatin regulatory factors, including TP53 and histone-modifying proteins, influence the accessibility of viral genomes and can regulate the transition between latency and productive replication by controlling chromatin state and transcriptional competence (23, 39, 92). In addition, cytoskeletal remodeling and cell adhesion pathways are essential for intracellular transport of viral components, nuclear egress, and cell-to-cell spread, processes that are extensively manipulated by alphaherpesviruses, including MDV (51, 111–113). Together, the functional coherence of these shared gene sets supports their likely mechanistic relevance in establishing a cellular environment permissive for efficient MDV lytic replication. Shared host DEGs clustered into proteostasis, immune modulation, transcriptional regulation, and cytoskeletal trafficking modules, suggesting a conserved host program supporting productive replication.

### Limitations and future directions

Some limitations of this study are worth noting. First, fully productive replication—characterized by efficient production of cell-free virus—occurs in epithelial skin cells for MDV and other alphaherpesviruses such as HSV and VZV. In contrast, the LCLs examined here are transformed CD4⁺ T-cell populations. Consequently, the regulatory architecture controlling reactivation in these cells is likely distinct from that in epithelial cells, which support high-level lytic replication and virus shedding *in vivo* (114–116). Second, while a temperature shift provides a reproducible and convenient trigger for MDV reactivation, it may also activate generalized cellular stress responses independent of virus-specific mechanisms (117, 118). We therefore prioritized candidate regulators that were consistently associated with viral transcriptional progression and host permissiveness across multiple contrasts and analytical approaches. Third, although analyzing two independent LCLs is more robust than the single-cell-line approach common in most studies, expanding the analysis to additional independent lines would further strengthen the findings. Finally, transcriptomic data alone cannot establish causal relationships. Functional validation through targeted gene expression, protein-level analyses, and genetic or pharmacological perturbation experiments will be essential to distinguish true regulatory drivers from downstream effects and to definitively identify host determinants of herpesvirus latency and reactivation.

### Proposed model of host-gated MDV reactivation

Despite these limitations, integration of viral transcription, host differential expression, pathway enrichment, and regulator analyses converged on five major biological domains that may govern permissiveness to MDV reactivation: chromatin accessibility and transcriptional competency, DNA damage and checkpoint signaling, proteostasis/unfolded protein response balance, mTOR-associated metabolic regulation, and innate immune gating. Collectively, these findings support a host-gated model in which cellular state determines whether the latent-to-lytic transition is initiated, whereas downstream productive replication proceeds through a comparatively conserved virus-driven program (Fig. 10).

**Fig 10.**
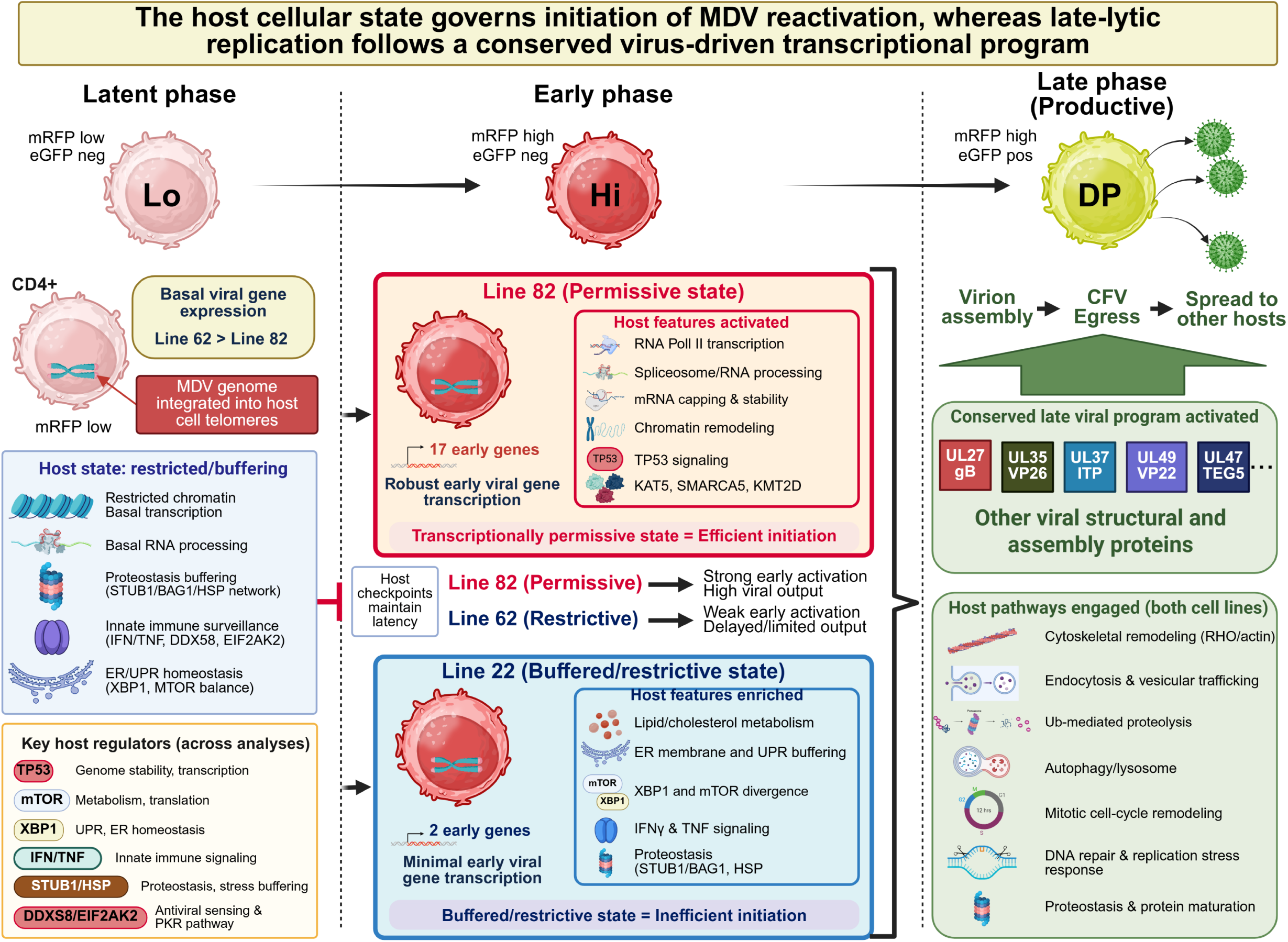
Proposed host-gated model of MDV reactivation in LCLs. During latency (Lo; mRFP low/eGFP⁻), the MDV genome is integrated in the telomeres of transformed CD4⁺ T cells, with minimal basal viral transcription. Line 62 had more basal viral gene transcription than Line 82. Host pathways associated with chromatin restriction, proteostasis buffering, innate immune surveillance, and endoplasmic reticulum (ER)/unfolded protein response (UPR) homeostasis contribute to maintenance of the latent state. Reactivation requires passage through a host-regulated checkpoint that determines whether cells enter productive viral replication. In the permissive Line 82 LCL, the transition from latency to early lytic replication (Lo to Hi) is accompanied by robust induction of 17 early viral genes, together with activation of RNA Polymerase II transcription, RNA processing, mRNA maturation, chromatin remodeling, and TP53-associated stress signaling. These transcriptional licensing programs establish a permissive cellular environment that supports efficient initiation of viral transcription. In contrast, the buffered/restrictive Line 62 LCL exhibited more basal viral gene expression during latency with limited early viral induction during reactivation. This was characterized by enrichment of lipid and cholesterol metabolism, ER-associated stress responses, IFNG/TNF signaling, altered MTOR and XBP1 activity, and proteostasis networks centered on STUB1/BAG1 and heat shock proteins, resulting in inefficient initiation of reactivation. Despite these early differences, both LCLs converge during the transition to productive late replication (DP; mRFP high/eGFP⁺), where a conserved viral transcriptional program dominated by structural and assembly genes is accompanied by coordinated host remodeling involving cytoskeletal reorganization, vesicular trafficking, ubiquitin-mediated proteolysis, autophagy, DNA repair, and protein maturation pathways. Together, the data support a model in which host cellular state governs entry into the MDV lytic cycle, whereas downstream productive replication proceeds through a largely conserved virus-driven transcriptional program.

### Summary

In this study, we differentiated MDV-reactivating LCLs into latent, early-lytic, and late-lytic phases of viral replication following temperature treatment, thereby enabling the identification of cellular and viral programs associated with the lytic switch. Importantly, studying two LCLs that maintain latency and reactivate differently is beneficial to our study, as it provides valuable information on how two LCLs, though they appear to be similar in overall reactivation competency, vary greatly in basal viral gene expression during latency and during the early phase of reactivation (lytic replication). Our findings demonstrate that MDV reactivation is strongly influenced by host cellular state. When comparing two LCLs in their responses to temperature-induced stress, clear differences are observed early during reactivation, but ultimately, both LCLs reach late lytic viral gene expression. Line 82 exhibits a transcriptionally permissive phenotype characterized by TP53-associated stress signaling, RNA biogenesis, and chromatin remodeling, while Line 62 displays a comparatively buffered state enriched for inflammatory signaling, altered metabolic regulation, and proteostasis-associated pathways. These differences predominantly influence the initiation of reactivation, while progression to late lytic replication is marked by a highly conserved viral transcriptional program accompanied by coordinated host remodeling of cytoskeletal, biosynthetic, and trafficking pathways.

Early induction of viral genes associated with nuclear egress and virion transport—including UL7, UL16, UL34, UL51, and US3—suggests that preparation for assembly and trafficking is a defining feature of the latency-to-lytic transition. Integration of host and viral analyses further identified TP53, MTOR, XBP1, NFKB1/RELA, STUB1, KAT5, SMARCA5, EIF2AK2, and DDX58 as candidate regulators that may govern permissiveness to MDV reactivation. Collectively, these findings support a model in which host regulatory architecture determines entry into productive replication, whereas downstream viral execution follows a comparatively conserved program. This work provides new insight into host–virus interactions that underlie MDV latency and reactivation and establishes a framework for future studies targeting cellular determinants of reactivation, with potential implications for improved control strategies and the development of next-generation vaccines against Marek’s disease and other cancer-causing herpesviruses.

## MATERIALS AND METHODS

### Ethics statement

All animal experimentation was conducted in accordance with national regulations. The animal care facilities and programs at the University of Illinois meet all requirements of the law (89–544, 91–579, 94–276) and NIH regulations on laboratory animals and comply with the Animal Welfare Act, PL 279. The University of Illinois Animal Care Program is accredited by the Association for Assessment and Accreditation of Laboratory Animal Care (AAALAC). All experimental procedures were conducted in compliance with approved Institutional Animal Care and Use Committee protocols. Water and food were provided to chickens ad libitum. Chickens were humanely euthanized using CO_2_ asphyxiation.

### Cell culture and cells

Chick embryo cells (CECs) were prepared from 10–11-day-old specific-pathogen-free (SPF) embryos obtained from the University of Illinois Poultry Farm using standard methods (119) and seeded in growth medium consisting of Medium 199 (Cellgro, Corning, NY, USA), 10% tryptose-phosphate broth (TPB), 0.63% NaHCO_3_, 100 U/ml penicillin, 100 µg/ml streptomycin, and 4% FBS (M24 media). Confluent CECs were maintained in Medium 199 supplemented with 7.5% TPB, 0.63% NaHCO_3_, antibiotics, and 0.2% FBS (M20.2 media). CECs were maintained at 38°C in a humid atmosphere of 5% CO_2_.

### MDV-induced lymphoblastoid cell lines (LCLs)

The recombinant (r)MDV used in this report has been previously described, named v2001 (17), in which the monomeric red fluorescent protein (mRFP) was fused to the MDV-specific early gene, RLORF4, termed RLORF4mRFP, and the enhanced green fluorescent protein (eGFP) gene was inserted in frame at the C-terminus of UL47, termed UL47eGFP. To generate LCLs, tumors were collected from Pure Colombian chickens infected with v2001 at four weeks post-infection. Single-cell suspensions were prepared by mashing tumor tissue through a 70 EASYstrainer (Greiner Bio-One, Monroe, NC, USA) in phosphate-buffered saline (PBS), and cell pellets were collected after centrifugation at 400 × *g* for 5 min at 4°C. Mononuclear cells were prepared using Histopaque 1077 (Sigma-Aldrich, St. Lois, MO, USA) by centrifugation at 400 × *g* for 15 min at 25°C. Purified mononuclear cells were cultured in LMH media [1:1 mixture of Leibovitz L-15 and McCoy 5A (LM) media (Gibco, Gaithersburg, MD, USA) supplemented with antibiotics (100 U/ml of penicillin and 100 μg/ml of streptomycin), with 10% FBS and 8% chicken serum] at 41°C with 5% CO_2_. After two weeks, the chicken serum concentration was gradually reduced to only 10% FBS (LM10 media). A total of four v2001 LCLs were established: Line 59, Line 62, Line 82, and Line 94. The LCLs were cultured for 20 weeks and used for subsequent experiments.

### Temperature-induced reactivation

LCLs were separated from cellular debris using Histopaque 1077 density gradient centrifugation as described above. Live cells were resuspended to 2×10^6^ cells/ml in LM10 and incubated at 41°C with 5% CO_2_ overnight. The next day, LCLs were temperature-treated by incubation at 32°C for varying durations. Temperature-treated cells were subsequently used for other studies.

### Cell-free virion extraction

To measure cell-free virions produced during reactivation, LCLs were temperature-treated, followed by cell-free virus extraction. After 24 h of temperature treatment, 10×10^6^ LCLs were collected by centrifugation at 200 × *g* for 5 min at 4°C, then resuspended in 1 ml of SPA buffer (7.5% sucrose in 0.01 M NaPO_4_ and 1.0% bovine serum albumin). Cell-free viral particles were prepared essentially as previously described (120). Briefly, LCLs were disrupted twice by sonication at ∼20 Hz for 10 s using a W-220 sonicator (Heat Systems-Ultrasonic, Inc., Farmingdale, NY, USA) on ice. Cell-free virions were collected by centrifuging at 1000 × *g* for 20 min to remove the debris. The supernatant was filtered through Millex AA 0.8 µm filters (Merck Millipore Ltd, Tullagreen Carrigtwohill, County Cork, Ireland) and used in plaque formation assays.

### Plaque formation assay with cell-free virions

Plaque-formation assays were used to quantify the number of LCLs that produced infectious cell-free MDV upon reactivation by temperature treatment. Briefly, cell-free virus extracts were serially diluted with M24 media and added to primary CECs seeded the day prior. The next day, the media was removed, and fresh M20.2 media was added. After 5-7 days, CECs were washed once with PBS, fixed/permeabilized with PFA buffer (2% paraformaldehyde, 0.05% Triton X-100 in 1× PBS) for 15 min at 25°C, and washed twice with PBS. Anti-MDV chicken sera and goat anti-chicken IgY-Alexa Fluor 488 or 568 secondary antibodies (Molecular Probes, Eugene, OR, USA) to detect MDV-induced plaques using an EVOS FL Cell Imaging System (Thermo Fisher Scientific, Waltham, MA, USA). Plaques were enumerated and expressed as plaque-forming units (PFU) per 1×10^6^ cells.

Statistical analyses of cell-free virus production were performed using IBM SPSS Statistics version 27 (IBM Corp., Armonk, NY, USA). Differences in PFU among experimental groups were evaluated using the Kruskal–Wallis test, a nonparametric alternative to one-way analysis of variance (ANOVA), followed by pairwise multiple-comparison testing when significant overall effects were detected. Statistical significance was declared at P < 0.05.

### Flow cytometry and cell sorting

For flow cytometry analysis, 8×10^6^ LCLs were harvested after temperature treatment as single-cell suspensions by filtering through a 70 µm EASYstrainer and diluted to 1×106 cells/ml before analysis on a Cytek Aurora spectral flow cytometer (Cytek Biosciences, Fremont, CA, USA). For measuring cell viability, 3 µM of DAPI (4’,6-Diamidino-2-Phenylindole, Dihydrochloride) staining solution was added to the single-cell suspension for 15 min before analysis by flow cytometry following the provided protocol by Invitrogen (Invitrogen, Carlsbad, CA, USA). The mRFP and eGFP data are shown after gating on lymphocytes, followed by single-cell and DAPI gating.

For cell type characterization, LCLs were stained with anti-chicken CD4-, CD8-, KUL01-, and Bu1-Alexa Fluor 647 antibodies (Southern Biotech, AL, USA) at 1:1000 dilutions. The data are shown after gating on lymphocytes, then applying single-cell, DAPI-negative, and mRFP gates. Ten thousand gated cells were acquired per sample and analyzed using FlowJo software (Ashland, OR, USA).

For cell sorting, cells were submitted to the Cytometry and Microscopy to Omics (CMtO) facility at the University of Illinois at Urbana-Champaign. The LCLs were kept at 4°C during cell sorting using a Bigfoot spectral cell sorter (ThermoFisher, USA). The LCLs were separated into three cell populations: mRFP low (Lo), mRFP high (Hi), and double-positive (DP) for cells expressing mRFP and eGFP. A total of 4×10^5^ cells were sorted for each population and used for RNA extraction and sequencing. The data were analyzed using FlowJo v10 software (BD Biosciences, San Diego, CA, USA), and population frequencies were calculated as percentages of live LCLs.

### Construction of stranded RNA-seq libraries

Total RNA was extracted from sorted 4×10^5^ cells using a Qiagen RNeasy kit with on-column DNase treatment, following the manufacturer’s instructions (Qiagen, Germantown, MD, USA). Stranded RNA-seq libraries were constructed at the DNA Services Laboratory of the Roy J. Carver Biotechnology Center at the University of Illinois at Urbana-Champaign. Total RNA was quantified using a Qubit Fluorometer (Life Technologies, Grand Island, NY, USA), followed by RNA Quality (RQ) analysis using an Agilent Bioanalyzer (Santa Clara, CA, USA). RNA samples with an RIN >9 were used for library construction. Forty nanograms of total RNA were used per sample to prepare libraries with the Ovation Universal RNA-seq kit from Tecan Genomics, Inc. (Morgan Hill, CA, USA). The final libraries were quantified using a Qubit Fluorometer, and the average size was determined using an AATI Fragment Analyzer (Advanced Analytics, Ames, IA, USA). Samples were diluted to 5 nM and further quantified by qPCR on a Bio-Rad CFX Connect Real-Time System (Bio-Rad Laboratories, Inc., CA, USA).

### RNA Sequencing (RNA-seq)

The final stranded RNA-seq library pool, consisting of 18 libraries, was sequenced on two SP lanes of an Illumina NovaSeq 6000 SP flow cell as single-end 100 nt reads. The run generated .bcl files, which were converted into adaptor-trimmed, demultiplexed fastq files using bcl2fastq v2.20 Conversion Software (Illumina, San Diego, CA, USA). The post-sequencing processes, including image analysis, base calling, and Q-Score calculation, were performed on CNRG’s Biocluster high-performance computing resource (http://biocluster.igb.illinois.edu/). All analyses, from the summation of counts to the gene level, were performed on a laptop in R version 4.2.1 (2022-06-23 curt).

### RNA-seq analysis, mapping, and gene expression quantitation

Sequencing data were processed by the High-Performance Biological Computing (HPCBio) team at the Roy J. Carver Biotechnology Center (University of Illinois at Urbana-Champaign). The chicken reference genome and transcriptome files (*Gallus gallus*; Annotation Release 106) were downloaded from the NCBI FTP site (https://ftp.ncbi.nlm.nih.gov/genomes/). Transcript quantification was performed using Salmon v1.5.2 (121), which implements a lightweight quasi-mapping approach to estimate transcript abundance relative to both the *Gallus gallus* reference transcriptome and the MDV strain RB1B genome (122). Transcript-level abundance estimates were summarized to gene-level counts using the tximport package in RStudio, applying bias-corrected counts without an offset method (Table S2). Raw gene counts were imported into the edgeR package for normalization and filtering.

To remove low-expressed genes, transcripts had to have at least ten counts in at least three samples (corresponding to the smallest replicate group). After filtering, 16,612 genes remained for downstream analysis, accounting for 99.93% of the total mapped reads. Library size and compositional biases were normalized using the trimmed mean of M-values (TMM) method. Following filtering, TMM normalization was repeated to ensure appropriate scaling of the retained gene set (123).

To account for the mean–variance relationship inherent to RNA-seq count data, the voom transformation from the limma package was applied to generate precision-weighted log2-transformed expression values suitable for linear modeling. To control for unwanted technical and biological variation, unwanted variation (RUV) factors were estimated using the RUV framework (124). One-way ANOVA across all six experimental groups was performed to evaluate the effects of increasing numbers of RUV factors. Based on stabilization of gene-level variance and model performance, five RUV factors were selected and incorporated into the final linear model. Differentially expressed genes (DEGs) analysis was performed using the limma-voom framework with the following model:

Expression ∼ Group + 5 RUV factors: Within each cell line, three pairwise contrasts were tested: Hi vs. Lo (reactivation/early lytic vs. latency), DP vs. Lo (late lytic vs. latency), and DP vs. Hi (late vs. early lytic progression). To ensure consistency in false discovery rate (FDR) estimation across contrasts, raw p-values from all six contrasts (three contrasts × two cell lines) were jointly adjusted using the Benjamini–Hochberg method (125). This global FDR correction ensured that identical raw p-values yielded identical adjusted FDR values across contrasts. Genes were considered differentially expressed at a primary threshold of FDR < 0.05. For exploratory systems-level analyses, including pathway and network inference, an additional relaxed threshold of FDR < 0.25 was applied. Complete statistical results for RNA-seq analyses are provided in Data S1.

### Functional enrichment and pathway analysis

To identify biological processes, signaling pathways, and regulatory networks associated with MDV reactivation, differential gene expression results were interrogated using Ingenuity Pathway Analysis (IPA, http://www.ingenuity.com/; Ingenuity Systems Inc., Redwood City, CA) and SRplot (https://www.bioinformatics.com.cn), an online platform for gene ontology (GO) and pathway enrichment visualization (20). For both enrichment platforms, the complete annotated RNA-seq dataset served as the background reference for statistical testing. Differentially expressed genes identified at FDR < 0.05 were used for primary high-confidence pathway inference. To explore coordinated but lower-amplitude transcriptional programs, additional analyses were performed on DEGs identified with an FDR < 0.25.

### Ingenuity Pathway Analysis (IPA)

DEGs identified using the limma-voom framework were imported into IPA for canonical pathway analysis, upstream regulator prediction, regulator effects modeling, and disease/biofunction annotation. Gallus gallus gene identifiers were directly mapped within the IPA knowledge base. Primary high-confidence pathway inference was performed using DEGs with FDR < 0.05. To identify broader regulatory trends and coordinated transcriptional programs, additional exploratory analyses were performed using DEGs defined at FDR < 0.25. Enrichment significance in IPA was determined using a right-tailed Fisher’s exact test. Activation states were predicted using the IPA activation z-score algorithm, where z ≥ 2 indicated predicted activation and z ≤ –2 indicated predicted inhibition. Canonical pathways and functional categories with P ≤ 0.001 were considered significantly overrepresented.

### Gene Ontology and KEGG Enrichment (SRplot)

Gene Ontology (GO) and KEGG pathway enrichment analyses were performed using the SRplot platform, as previously described (21). Importantly, SRplot analyses were conducted exclusively using DEGs identified at FDR < 0.25. This relaxed threshold was selected to capture coordinated systems-level transcriptional patterns that may not reach stringent gene-level significance but represent biologically coherent pathway-level changes. Gene lists from each contrast (Hi vs Lo, DP vs Hi, and DP vs Lo) were analyzed separately for each cell line. Enrichment testing was performed using a hypergeometric test, and multiple-testing correction was applied using the Benjamini–Hochberg method. Enriched GO terms and KEGG pathways with adjusted P-value (FDR) ≤ 0.05 were considered statistically significant. SRplot was used to generate dot plots, bar plots, and GO three-ontology visualizations representing gene ratio, gene counts, and adjusted enrichment P-values. All enrichment results, including canonical pathways, upstream regulators, and regulator networks, are provided in Data S2.

### Integration of Functional Enrichment Approaches

Together, IPA and SRplot provided complementary and mechanistically distinct functional perspectives on the transcriptional changes associated with MDV reactivation. SRplot-based GO and KEGG enrichment analyses provided an unbiased, gene-set–centric overview of overrepresented biological processes, molecular functions, and pathways, thereby quantifying the global functional landscape of differential gene expression for each contrast and cell line. In contrast, IPA enabled regulator-centric modeling, incorporating curated molecular interaction networks to predict upstream regulators, infer activation states (z-scores), and construct causal networks linking host transcriptional programs to viral reactivation states. This combined analytical strategy allowed hierarchical interpretation of the data at multiple confidence levels. Specifically, it enabled identification of: 1) High-confidence pathways derived from DEGs meeting stringent thresholds (FDR < 0.05), supporting robust, regulator-supported biological conclusions. 2) Coordinated systems-level trends were identified using relaxed thresholds (FDR < 0.25), capturing lower-amplitude but biologically coherent transcriptional programs relevant to lytic progression. 3) Shared versus cell-line–specific regulatory modules, permitting direct comparison of transcriptional permissiveness between Lines 62 and 82. 4) Candidate host determinants of MDV lytic permissiveness, identified through convergence of canonical pathways, upstream regulator predictions, and network architecture.

By integrating enrichment-based overrepresentation analysis (SRplot) with causal network inference (IPA), the study distinguished between broad transcriptional remodeling, regulatory gating mechanisms, and conserved viral–host interaction modules. This layered approach strengthens the mechanistic interpretation of host-dependent differences in MDV reactivation and provides a systems-level framework for identifying host factors that govern entry into the lytic cycle.

## ACKNOWLEDGEMENTS

The authors would like to thank Nagendraprabhu Ponnuraj for help during animal experiments and for collecting samples. The authors would also like to thank Jenny Drnevich in the High Performance Computing in Biology core at the Roy J. Carver Biotechnology Center for her help in performing and analyzing RNA-seq data. This work made use of the equipment, software, and facilities provided by the University of Illinois Urbana-Champaign College of Veterinary Medicine Shared Equipment Program’s Biocomputing Shared Resource (BioShaRe). The College of Veterinary Medicine BioShaRe is housed in the Illinois Campus Cluster, a computing resource operated by the Illinois Campus Cluster Program (ICCP) in conjunction with the National Center for Supercomputing Applications (NCSA), and supported by funds from the University of Illinois at Urbana-Champaign.

## Funding

This report was supported by a National Institute of Food and Agriculture Hatch grant ILLU-888-936 from the USDA National Institute of Food and Agriculture for K.W.J. Dr. Yung-Tien Tien was supported by a Taiwanese University of Illinois at Urbana-Champaign Scholarship.

## Author contributions

Conceptualization: Y.T., K.W.J.

Data curation: H.A., Y.T., K.V.E.

Formal analysis: H.A, Y.T.

Funding acquisition: K.W.J.

Investigation: H.A., Y.T., K.V.E., K.W.J.

Methodology: Y.T., K.V.E., K.W.J.

Project administration: K.W.J.

Visualization: H.A., Y.T., K.W.J

Supervision: K.W.J.

Writing—original draft: H.A., Y.T.

Writing—review & editing: H.A., K.W.J.

## Competing interests

The authors declare they have no competing interests.

## Data, code, and materials availability

All data and code needed to evaluate and reproduce the results are present in the paper and/or the Supplementary Materials. LCLs used in this article are available upon request. Raw transcriptome sequencing data are available in NCBI under the Gene Expression Omnibus Accession GSE338813.

